# Concurrent ecological and evolutionary processes contribute to mutualism breakdown between legumes and rhizobia

**DOI:** 10.64898/2025.12.02.691918

**Authors:** Sierra L Bedwell, Isabelle M Lakis, Allison R Megow, Angelina E Carpenter, Kevin D Ricks, Christopher J Fields, Jennifer A Lau, Rachel J Whitaker, Katy D Heath

## Abstract

Ecology and evolution jointly shape community responses to environmental perturbations, yet these processes are often examined separately, even in microorganisms where both occur over short time-scales. For example, during mutualism breakdown, shifts in diverse microbial communities might coincide with evolutionary changes in mutualistic traits within key symbionts. Here we study the legume-rhizobium mutualism after 33 years of nitrogen fertilization. Pairing a manipulative inoculation study with full-length 16S rRNA amplicon sequencing provides incomparable resolution across biological scales: whole bacterial community, genus *Rhizobium, Rhizobium* ASVs, and symbiosis plasmids. Clover host plant growth is first limited by symbiont availability; clover’s preferred partner declines in N-addition soils, while a diverse, largely uncharacterized *Rhizobium* community increases. This ecological change is compounded by a concurrent evolutionary decline in the quality of the preferred symbiont through changing frequencies of symbiotic plasmids. These shifts make mutualists both rarer and inferior, reducing the benefits hosts receive from mutualism.

## Introduction

Microbial mutualisms are some of the most important mutualisms on Earth, altering biogeochemical cycles^1,2^, promoting plant productivity^3,4^ and protecting hosts from stress and enemies^5–7^. These mutually beneficial interactions between species are fundamental to every ecosystem and yet may be unstable^8–12^. Under environmental change, mutualisms may break down, with partners potentially evolving to parasitize their host, switching to new hosts, or adopting a free-living lifestyle^12–14^. Thus, understanding why and how mutualisms break down will be essential to maintaining them in the face of a changing world, and tractable mutualisms offer ideal models with which to examine the interplay of ecology and evolution in this process.

Mutualism breakdown has been observed in multiple systems due to either ecological or evolutionary changes. Ecological changes such as community-level shifts in composition can drive decline; for example, multiple studies have found that invading ants can disrupt the existing mutualism between African ants and acacia trees^15,16^. Additionally, breakdown through evolutionary mechanisms such as population-level shifts in allele frequency has been observed in diverse systems such as corals and pollinator wasps^12,17–19^. Though models have predicted that mutualisms can break down through concurrent ecological and evolutionary processes^20,21^, this has been difficult to measure in natural systems. In macroorganisms, generation times are slow enough that incorporating evolutionary processes into short-term experimental systems is difficult, while in microbes, ecology and evolution occur on comparable time-scales, yet cannot easily be differentiated^22,23^.

In microbial communities, allele-frequency shifts within populations of single microbial species may occur alongside community composition shifts at broader taxonomic scales^23^. Yet, in microbes, these processes have often been examined using different techniques. For example, population genomics and classical experimental evolution provide high-resolution of allelic shifts within species but: 1) require isolates be culturable^24^ and 2) may miss key dynamics occurring between species in a more complex community^25,26^. In contrast, standard amplicon sequencing (e.g., the V4 region of the 16S rRNA) can capture a large portion of unculturable diversity in complex microbial communities, but has been limited by amplicon length and sequencing depth, thus restricting fine-scale resolution to move beyond community shifts at the broad scale (phylum, order, and family)^27,28^. Additionally, microbial populations and communities are interconnected by the movement of genes within and between taxa through horizontal gene transfer (HGT)^29,30^. For microbial mutualists with mobile symbiosis genes^31,32^, this movement may accelerate both the emergence and breakdown of mutualism^33–35^, as well as potentially blurring the lines between ecological and evolutionary processes^23,36^.

The legume-rhizobium mutualism is an ideal model to examine the ecological and evolutionary dynamics of mutualism breakdown because it consists of a taxonomically limited set of focal symbionts within progressively more complex communities, from the internal structures of the host plant to the hyperdiverse soil environment. In the resource mutualism between leguminous plants and nitrogen-fixing rhizobia, rhizobial cells living in soil form nodules on legume roots and therein fix atmospheric dinitrogen gas into plant-available forms of nitrogen (N) in exchange for carbon from the plant host^4,37^. In the *Trifolium* (clover)*-Rhizobium* symbiosis, and many others, the mutualism relies on the presence of symbiosis genes carried on mobile genetic elements (MGEs), often plasmids (pSyms) or integrative conjugative elements (symICEs)^32,34^.

Previous work in this system has shown that mutualism can break down under long-term fertilizer (N) addition, and culture-dependent studies suggested an evolutionary explanation by identifying genetic changes located within the pSyms that differentiated isolates from fertilized versus unfertilized populations^38,39^. In fact, isolates collected from N-addition soils often contained evolutionarily distinct pSyms that conferred worse partner quality^40^. However, when these genetic effects were isolated using single inoculation experiments, the impacts on host growth were much smaller compared to those estimated when plants were inoculated with whole soil from those same plots^38^, suggesting that ecological changes in rhizobial community composition and/or abundances, in addition to rhizobium evolution, also play a key role in mutualism breakdown between clover and rhizobia under long-term N-addition^38–40^.

Addressing potential ecological changes that may have taken place in these communities under long-term fertilization requires a much deeper understanding of this extremely diverse group (the genus *Rhizobium* as well as the larger family *Rhizobiaceae*). This group has posed taxonomic challenges – both because naming conventions for plant symbionts often predate modern microbial taxonomy^41^ and because, until recently, there were insufficient full reference genomes distributed across the entire family with which to discriminate chromosomal genera^42,43^. The family *Rhizobiaceae* consists of at least 38 genera and nearly 300 species, which are linked by extensive HGT and plasmid exchange^42,44,45^. Yet even this vast rhizobial diversity comes mainly from nodule isolates, which may have experienced significant host filtering and may not represent overall community diversity^46–48^. For example, our previous culture collection of 63 isolates only captured two genospecies (gsE and gsB^38^) – only 11% of the total *R. leguminosarum* genospecies diversity identified in clover nodules worldwide^43^. Culture-dependent work has suggested that rhizobia in soil are rare compared to other bacterial taxa, yet may be diverse^49,50^. Thus making sense of rhizobial communities using standard culture-independent methods is challenging, as short-read amplicon sequencing often lacks both the resolution to differentiate disparate genera within the family^43,44^ and the depth to capture rare taxa in hyperdiverse communities. It is clear that understanding any ecological drivers of breakdown within this mutualism requires high-resolution, culture-independent data that can characterize rhizobial diversity in and out of nodules.

Here we leverage a long-term N-fertilization experiment with high-throughput, long-read amplicon sequencing of microbial communities from soil, rhizosphere, and nodules from a common garden greenhouse inoculation experiment to determine how 33 years of N-fertilization has affected legume-associated microbial mutualists at multiple scales. First, we examined the diversity of microbial communities across compartments. Then, we focused on ecological community shifts of rhizobial taxa, as well as evolutionary changes in a conserved pSym gene, *nodA*. Here, we model how these concurrent ecological and evolutionary forces contribute to a decline in mutualistic partner quality and thus clover host biomass. Finally, we examine the phylogenetic diversity of the whole rhizobial community throughout plant compartments and between N-addition and control, to determine how this important symbiont community responds to environmental change. In this work we unpack how the microbial community, across all levels, is impacted by N-fertilization across plant compartments, from field soil, to greenhouse soil, to plant rhizospheres, to symbiotic nodules.

## Methods

Full details for all experiments and analyses, as well as DOI links to our Zenodo repository, are provided in Supplementary Data. We summarize the key approaches below. All data are available on Zenodo.

### Study site and soil collection

Our inoculum comes from the successional long-term N-fertilization experiment (the T7 untilled microplots) at the Kellogg Biological Station Long-Term Ecological Research Site (KBS LTER) in Michigan, USA. This experiment includes six pairs of 25 m^2^ plots: with one plot in each pair fertilized since 1989 with agriculturally relevant levels of nitrogen and the other remaining unfertilized^51^. Three clover species have been recorded in the unfertilized plots since their inception, but have been almost absent from the N-addition plots since the 1990s^51–53^. We sampled field soil in July 2022 after 33 years of N-fertilization; taking 48 soil cores in total (4 per plot x 6 plots per N treatment x 2 N treatments).

### Greenhouse experiment

We grew plants in the University of Illinois Greenhouse. We inoculated 2 mL slurries from each core onto separate pots, containing a single one-week old seedling of either one of two clover species-*Trifolium hybridum* and *T. repens*. Inoculants and clover species were randomly assigned to pots in each of three replicate blocks. Plants were never provided exogenous nitrogen; this was to ensure that the only N the plants received throughout the course of the experiment came from microbial production from the microbes contained within the slurries.

Each block contained 126 plant pots including 30 controls (16 plants grown in autoclaved soil, 10 plants inoculated with sterile PBS, and 4 no-plant pots inoculated with live soil). Plants were harvested after 8 weeks. We separated aboveground and belowground biomass (belowground tissue was placed into a tube of sterile 1X PBS), and additionally collected one 15 mL tube of bulk soil from each pot. In the lab, we weighed aboveground biomass for all plants after drying for x days at 60°C. Belowground samples were split by block, 1: destructive nodule counting, 2: DNA sequencing, and 3: Long-term storage. For sequencing, we separately processed bulk soil, rhizosphere, and nodule samples.

### PacBio Revio full-length 16S rRNA and nodA sequencing and sequence processing

We sequenced full-length 16S rRNA from source soil, bulk soil, rhizosphere, and nodule samples, and *nodA* from the nodules. To determine whether or not relative abundance could be used as a proxy for absolute cell counts, we sequenced an additional 10 bulk soil samples by adding 20 μL of a commercial spike-in to the 500 mg soil sample prior to DNA extraction (ZymoBIOMICS Spike-in Control II Low Microbial Load, Zymo Research, CA, USA).

Library construction and sequencing on the PacBio Revio (Pacific Biosciences, CA, USA) were performed at the Roy J. Carver Biotechnology Center, University of Illinois.

Initial sequence processing was completed by the High Performance Computing Group (HPCBio) at the University of Illinois. To process our 10 bulk soil absolute abundance samples, we employed a custom R script to quantify absolute cell counts and determine that cell counts were correlated with relative abundance for *Rhizobium* and other randomly selected genera (Fig. S1).

After quality filtering and post-hoc chimera removal, we retained 7761 ASVs across 389 genera. Saturating rarefaction curves indicated that even after filtering we captured the majority of diversity present (Fig. S2).

### Rhizobium classification

To focus on clover symbionts, we used the QIIME delineation of the genus *Allorhizobium-Neorhizobium-Pararhizobium-Rhizobium* (referred to hereafter as the ANPR group) for initial classification (172 ASVs). To classify these ASVs below the genus level, while accounting for the extreme polyphyletic nature of the *Rhizobium* genus, we used the updated *Rhizobiaceae* framework^42^, placing references alongside ASVs in an

IQ-TREE phylogeny, and BLASTing some key ASVS prior to building the tree. Full-length 16S rRNA resolves the clade comprising *Rhizobium* sensu stricto (*leguminosarum*–*etli*), *Martinezella* (*tropici*–*rhizogenes*), and *Arminella* (*tubonense*–*tumorigenes*) from more distant genera, but does not reliably differentiate these three genera from one another; we therefore refer to this clade as *Rhizobium* sensu lato.

Forty-four ASVs were assigned to *Rhizobium* sensu lato on this basis. To more specifically classify genospecies members of the *Rhizobium leguminosarum* species complex (*Rlc),* we downloaded a reference database^43^. Rarefaction curves of the genus revealed that these ASVs represented the majority of the diversity of this genus, both in environments where *Rhizobium* was at a very low abundance (source soil) and at a very high abundance (nodules) (Fig. S2).

### Usage of large language models (LLMs)

To generate and edit code, as well as to edit text for grammar, the LLMs ChatGPT (OpenAI, CA, USA) and Claude (Anthropic, CA, USA) were utilized. The authors then reviewed and edited all LLM-produced content and take full responsibility for the content of this publication.

## Results

### The microbial community was strongly structured by plant compartment

As expected, the largest source of variation in microbial community composition was plant compartment (*i.e*., source soil (directly from the field), bulk soil (from pots in the greenhouse), rhizosphere (directly in contact with plants), or nodule), explaining 32% of total variance (Table S1A, S1B and Fig. S3). Inoculum source (whether the microbial community came from control versus N-addition soil slurries) and the compartment x inoculum interaction contributed 1.2% and 3.7% of the total variance overall, respectively. However the effects of inoculum source varied considerably, in within compartment-analysis, inoculum source effects were strongest in the field source soil (31.8% of variance), weaker in the greenhouse bulk soil and rhizosphere (3.4 and 3.5% of variance respectively), and not significant in the clover nodule (Table S1B). Whereas the source (field) and greenhouse bulk soil were dominated by common soil genera (*e.g*., *Bacillus*, *Nocardiodes,* Supplementary Data (SD) 1-3), the rhizosphere and nodules harbored plant-associated communities including the symbiont genera *Allorhizobium-Neorhizobium-Pararhizobium-Rhizobium* (ANPR). (SD1-3).

### Nodules are diverse yet dominated by a few ANPR ASVs

Because nodule taxa might be important drivers of clover health, we first asked which genera and ASVs dominated the nodule compartment. Nine different genera had mean relative abundances of above 1% in nodules (Fig. 1A), including many known non-rhizobial endophytes (NREs) such as *Cupriavidus, Pseudomonas,* and *Roseateles.* 20% of nodules overall were dominated by one of these non-rhizobium taxa (Fig. S4). Nonetheless, ANPR was the most abundant genus in clover nodules, with a mean relative abundance of 58%, a pattern which did not differ between fertilized and unfertilized inoculum sources (Fisher’s exact *P* value = 0.1674). While we had limited power to interrogate the impacts of individual NREs on plant biomass, we found no difference in plant biomass between samples dominated by ANPR versus other NREs, likely because even when nodules were not dominated by ANPR, half (11/22) maintained at least a 25% relative abundance of ANPR (results not shown).

**Figure 1:**
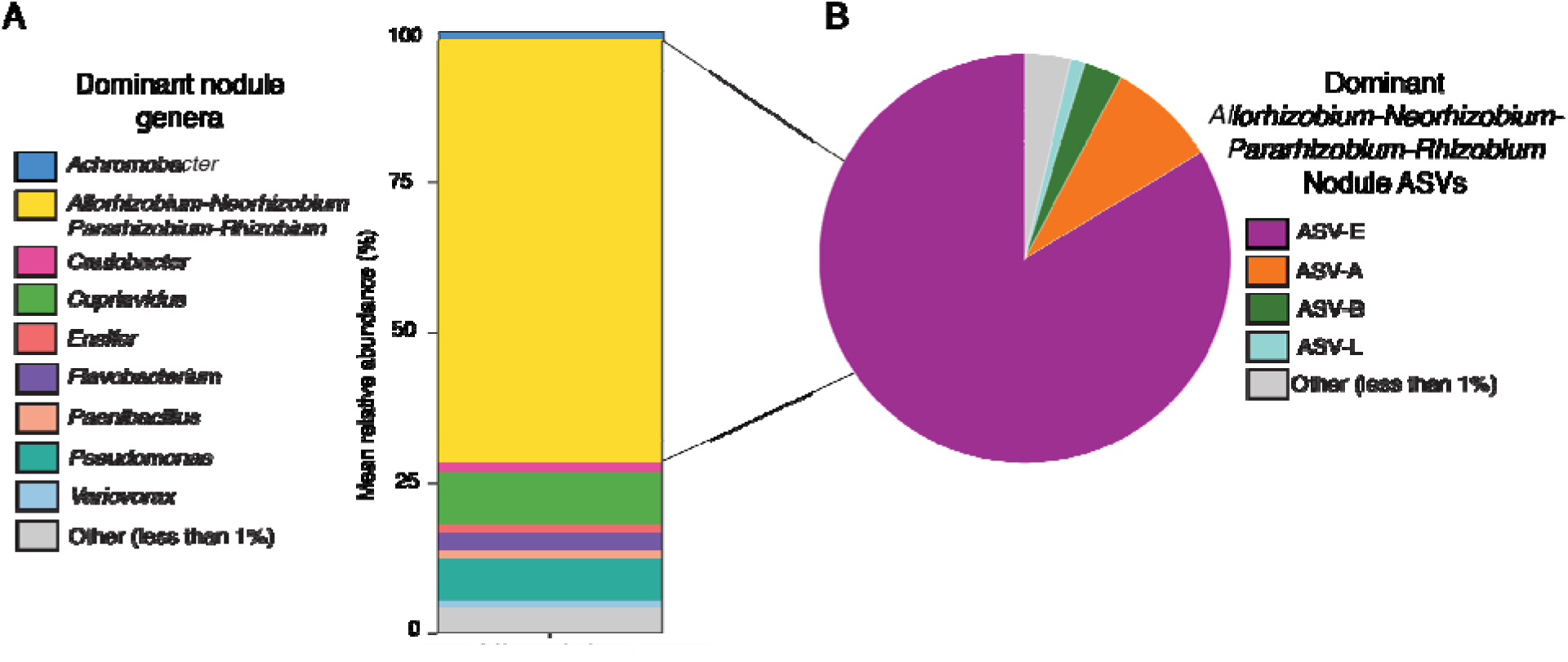
Nodules are diverse yet dominated by a few *ANPR* ASVs. **A**: Composition of nodules at the genus level between N and C. **B:** Composition of *Rhizobium* ASVs between N and C. For both, stacked bars represent the mean relative abundance of genera or ASVs between nodule samples. Each individual sample (comprised of 10 nodules per plant homogenized) was relativized, then all relative abundances were averaged. All genera/ASVs comprising less than 1% of reads in individual nodule samples were collapsed to “Other” (represented in gray and not present in the overall figure legend).

Among the ANPR ASVs in the nodules, only four maintained an abundance of more than 1% (Fig. 1B). To identify these ASVs, we BLASTed them against an up to date *Rhizobiaceae* database (SD8-9)^42^. The most abundant (ASV-E) was identical to the 16S rRNA sequence for *R. leguminosarum sensu stricto* (gsE), which was previously found to dominate clover nodules in this environment^40^; this sequence also matches several other genospecies (such as *R. brockwellii*/gsA*, R. laguerreae*/gsR*, R. ruizarguesonis*/gsC) and the more distant *R. hidalgonense* – none of which have been isolated from nodules from KBS. The ANPR taxon ASV-E was dominant within nodules (Fig. 1B), and was the second most ubiquitous ASV in the whole microbial community, present in 93% of all samples (SD4-6). Three other ANPR ASVs had mean relative abundances above 1% in nodules: ASV-B (an ASV which shared 100% identity to *R. johnstonii* (gsB) and *R. redzepovicii),* ASV-A (100% identical to *Agrobacterium arsenijevicii* and *A.* tumefaciens) and ASV-L (which was a single SNP different from *R. altiplani*).

### ASV-E is both less abundant and lower quality under N-addition

Given the dominance of ASV-E, as well as previous work implicating gsE’s role in clover health in this system^38,40^, we next examined how the relative abundance of ASV-E (relative to other ANPR ASVs) changed between N-addition and control inocula, and how these shifts related to plant growth in our inoculation study. We found that ASV-E was ∼3-fold less abundant in the N-addition source (field) soils (p < 0.0001, Fig. 2A, Table S2) and ∼1.7-fold less abundant in bulk soil of plants inoculated with N-addition field soil slurries (p < 0.05, Fig. 2A, Table S2), though there was no difference within the rhizosphere or nodule compartments.

**Figure 2:**
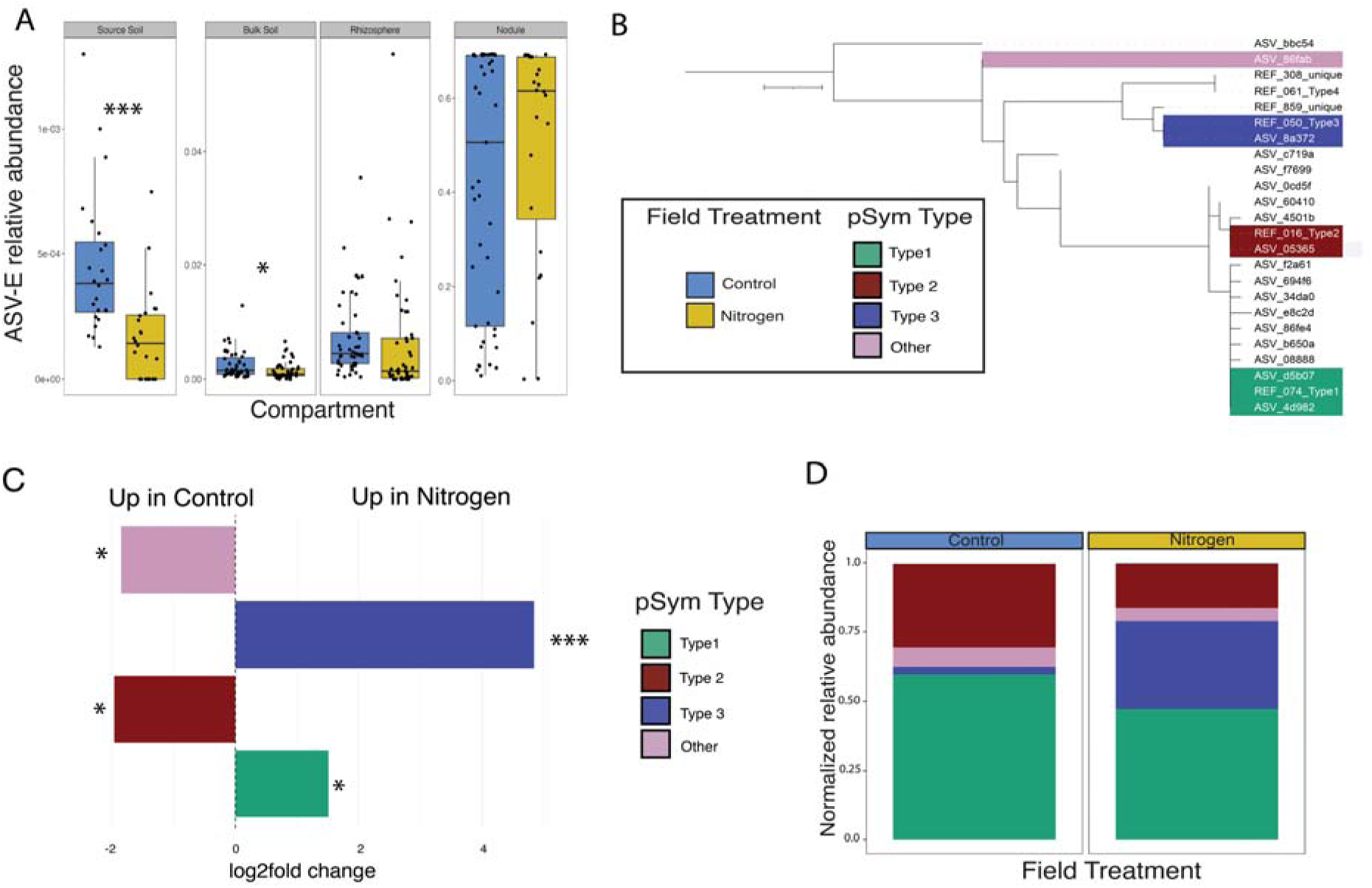
ASV-E is both less abundant (A) and dominated by low-quality pSym (B,C) under long-term N-addition. A: Relative abundance of ASV-E is significantly lower (p < 0.05) in nitrogen field source and bulk soil. Significance determined using linear mixed model considering nitrogen treatment and plot (nested within N-treatment). Compartments were modeled individually due to the large fold-changes in relative abundance between compartments which masked nitrogen effects. **B: nodA phylogenetic tree.** Reference sequences from our recent study^42^. Tree computed using MAFFT/IQTREE and visualized in ggTreeExtra. **C: log2fc of nodA**. Significance determined using DESeq2, *P* for Clade 3 = 1.0 x 10^-11^, *P* for “other” = 2.5 x 10^-2^. **D: nodA stacked bar plot**. Stacked bar plot depicts relative abundance normalized by sample number (Control = 45, Nitrogen = 24). Within B, C, and D, the two type 1 ASVs contained one SNP toward the end of their sequenced region which was ultimately trimmed out so all ASVs would be the same length as the reference genes; thus, the reads of these ASVs were combined in all analyses done on Type 1.

We also quantified the relative frequencies of previously characterized^40^ high-quality and low-quality pSym types in nodules from N-addition and control inocula using a marker of pSym type (*nodA*). An ASV matching to low-quality pSym type 3 was >28-fold enriched in nodules formed by plants inoculated with N-addition field soil slurries (Fig. 2B, Fig. 2C). Additionally, we found three more dominant *nodA* types (together, all four representing over 98% of sequencing reads). One ASV matched 100% to high-quality nodA type 2 (Fig. 2B), while two additional ASVs, one very common and the other extremely rare, both matched 100% to high quality nodA type 1. Additionally, a third abundant type (ASV_86f) has not yet been described, yet was more closely related to our *R. leguminosarum nodA* sequences than our natural outgroup (a rare *Rhizobium nodA* ASV with over 20 SNPs to all other ASVs; Fig. 3B). This ASV, though the rarest of the dominant four, was slightly enriched in nodules plants inoculated with control (versus N-addition) soil slurries (Fig. 2C, Fig. 2D).

**Figure 3:**
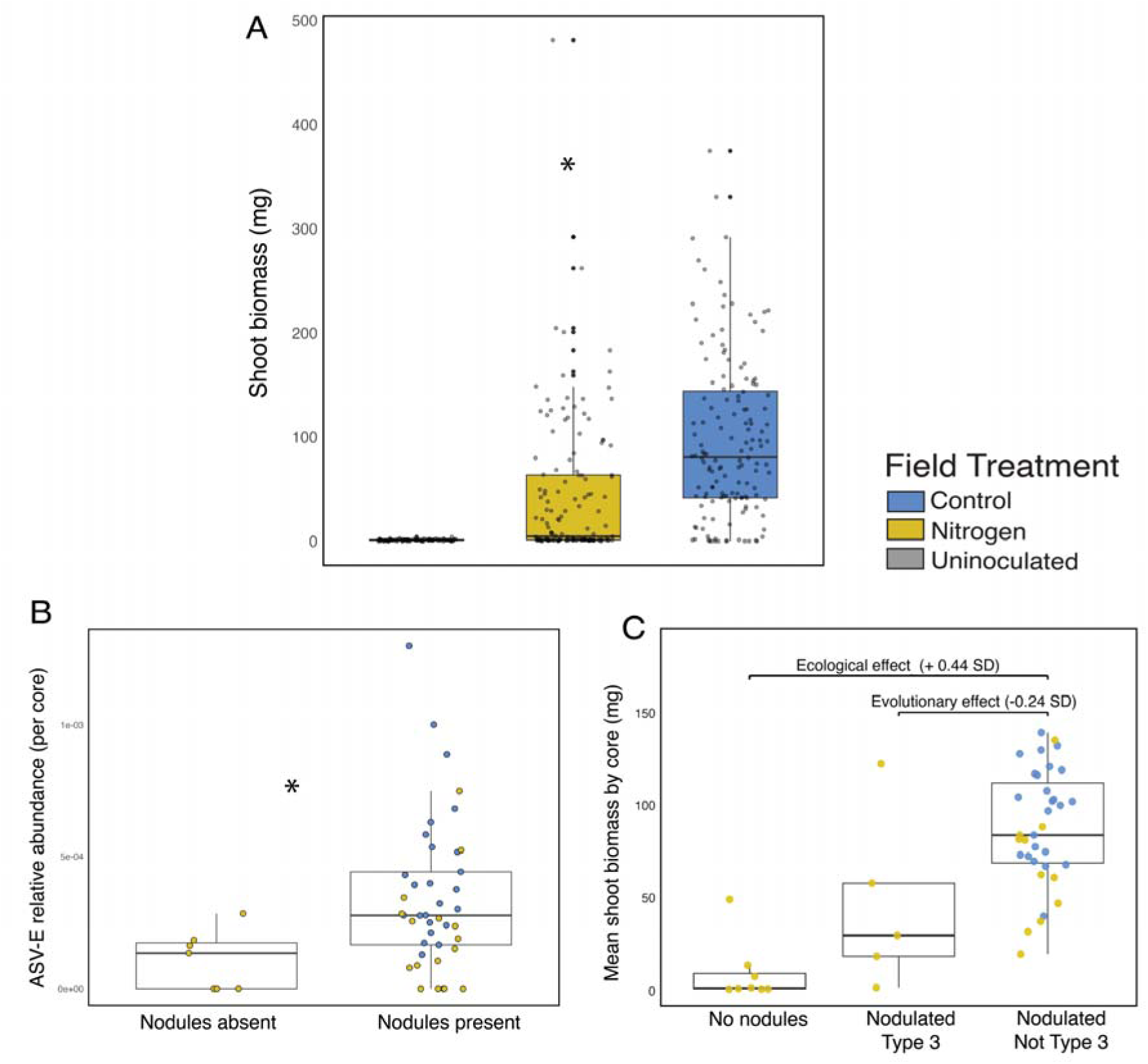
ASV-E relative abundance and pSym type predict plant health. A: Aboveground plant biomass is significantly lower when plants are grown in soil from N-deposition plots (F_1,284_ = 22.0, *P* < 0.01, N = 288h).**B: Relative abundance of ASV-E in the source soil predicts whether clover nodulates.** N = 48. Significance was determined using a Wilcoxon rank-sum test (Mann-Whitney U Test), *P* = 0.017. **C: Regardless of N-fertilization, when pSym type 3 is dominant in nodules, plant biomass is significantly lower.** Dominance was defined as over 50% of the relative abundance of *nodA* within a nodule being comprised of pSym type 3. Ecological and evolutionary effect sizes come from a two-part piecewise structural equation model; the ecological effect takes into account both the direct impact of ASV-E in source soil on biomass as well as the indirect effect of ASV-E ➔ nodulation ➔ biomass, the evolutionary impact takes into account the direct effect of nodules being dominated by the Type 3 pSym on biomass.

These observed changes in soil microbial communities were associated with differences in plant growth. Plants inoculated with soil slurries from long-term N-fertilization treatments produced 55% less biomass (Fig. 3A). We fit a two-part piecewise structural equation model (SEM) across all 48 source soil cores to partition the standardized effects of ecology (the direct and indirect, nodulation-mediated effects of ASV-E relative abundance on biomass) from evolution (the direct effect of type 3 pSym dominance on biomass). The ecological and evolutionary impacts did not differ in magnitude (bootstrap *P* = 0.33), with the total standardized ecological effect on biomass 0.44 (direct + nodulation-mediated indirect paths), and standardized evolutionary effect of the pSym type associated with a 0.24 SD decrease in biomass (p = 0.069). Within the ecological path, the indirect path (effect size 0.33) contributed more than the direct path (effect size = 0.11), suggesting that the majority of ASV-E’s ecological impact on plant biomass flows through nodule formation, rather than a direct impact. This is additionally supported as both legs of the indirect path were significant (ASV-E → nodulation, *P* = 0.038; nodulation → biomass, *P* = 0.002), whereas the direct ASV-E → biomass path was not (p = 0.441). All coefficients can be found in Table S3.

### Most ANPR ASVs are not Rhizobium, and few are compartment specialists

Next we wanted to examine rhizobial community composition and changes under N-addition more deeply, we classified the 172 ANPR ASVs beyond the SILVA delineation (*Allorhizobium-Neorhizobium-Pararhizobium-Rhizobium*). We constructed a phylogeny by using up-to-date references across the family *Rhizobiaceae* (SD7-8)^42^. From this tree, we were able to separate out 44 ASVs (26%) in the genus *Rhizobium* sensu lato, (three genera formerly known as *Rhizobium* and just recently split)^42^ which cleanly separated from the rest of the genera within the family (Fig. 4). We also observed that, contrary to our expectations by filtering our initial selection to the ANPR genera, the majority of our ASVs (54%), did not group with any of the four ANPR genera, and instead were most closely related to ∼10 additional genera within the family *Rhizobiaceae* (Fig. 4). Though many of the genera we observed have only been defined after the initial identification database we used was released (such as *Onobrychicola,* described in 2022, of which we had 18 ASVs), we identified one ASV which, at least by 16S rRNA, was phylogenetically similar to *Shinella,* an entirely separate genus in SILVA that we had explicitly filtered out in our analyses (Fig. 4). Another ASV, which despite passing all our filtering cutoffs, had less than 95% identity to every reference and a branch length 300 times longer than its neighbors, so was subsequently removed from our phylogenetic tree. Additionally, the majority of these ASVs (80% percent) had no perfect matches to known *Rhizobiaceae* and likely represent a significant expansion of known diversity in the family. Both the larger ANPR group, nor the true *Rhizobium* senso lato genus were compartment generalists, with 70% (122/172), present in both plant-associated (rhizosphere/nodules) and soil (bulk/source) environments. Additionally, though nodule presence was not common (only 27% of ANPR ASVs were seen in nodules), it was not phylogenetically restricted to the true *Rhizobium* genus (Fig 4).

**Figure 4:**
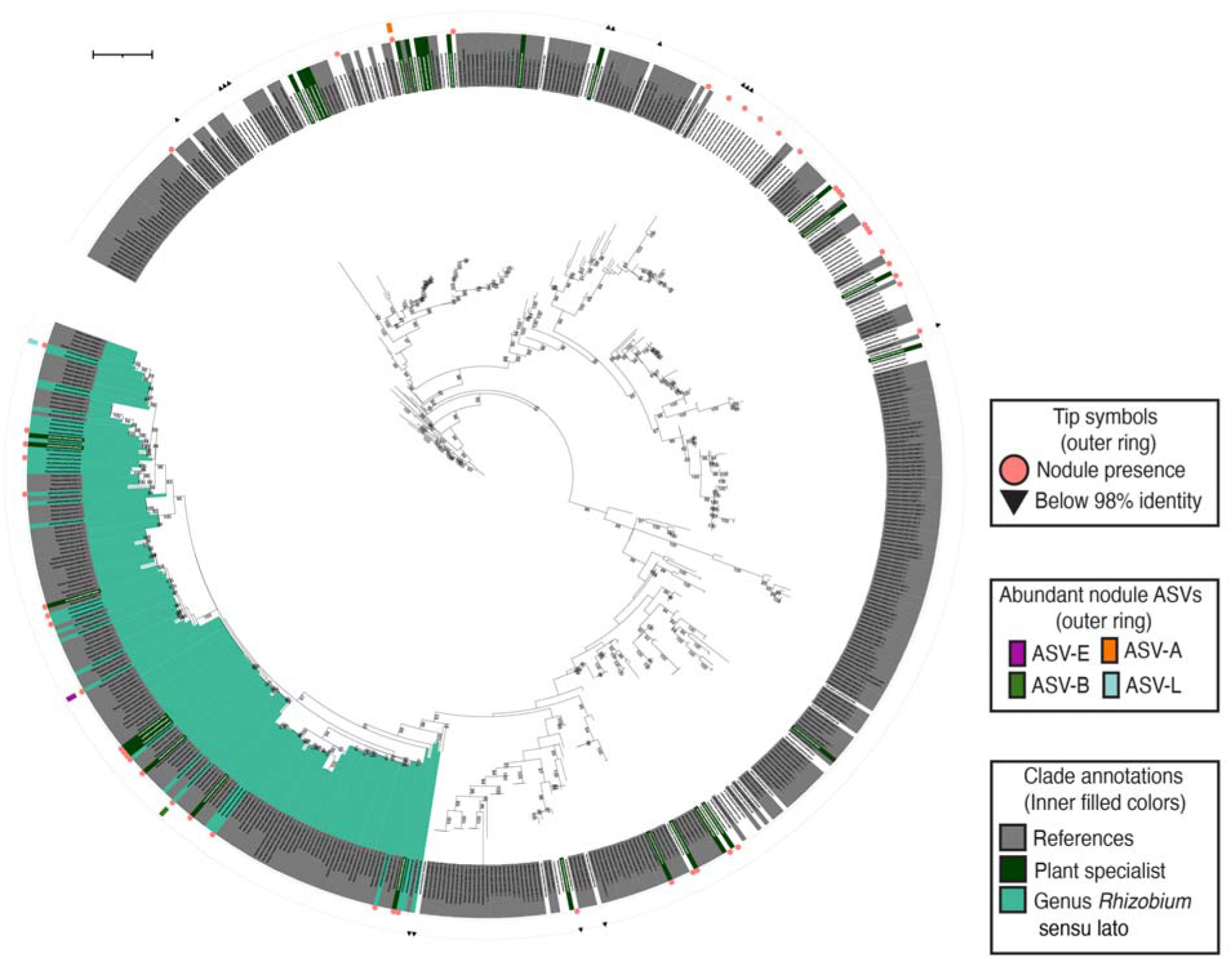
ASVs filtered to Allo-Neo-Para-Rhizobium span the entire family *Rhizobiaceae* and contain 44 *Rhizobium sensu lato* ASVs. Phylogenetic tree of 174 *Allorhizobium-Neorhizobium-Pararhizobium-Rhizobium* ASVs (one ASV was removed due to extremely low phylogenetic relatedness to everything else) and 245 reference sequences (labeled in grey) from the family *Rhizobiaceae*^42^. Tree is rooted on the genus *Mesorhizobium*. The three genera classified as *Rhizobium* sensu lato^42^ are not resolvable at the level of the 16S rRNA gene and are colored in teal. Plant specialist ASVs (only found in rhizosphere and/or nodules) are labeled in green. Any presence in nodules is recorded as a red circle on the outside. Abundant nodule ASVs (above 1% relative abundance overall) are labeled on the outside as colored rectangles. References are labeled in grey and ASVs that are only found in plant compartments (rhizosphere and nodules) and never soil are labeled in purple. Genera within *Rhizobiaceae* are labeled in colors within clades (and extending to branches for ASVs that are not plant specialists or references). Genus delineations, including genus *Rhizobium*, were defined from previous work^42^. Scale bar represents 0.1. A tree with all genera labeled can be found on iTOL; link in the Supplementary Methods.

### Genus-level aggregation masks ASV-level responses to N-addition

Using this refined taxonomy, we focused on the *Rhizobium* sensu lato genus to understand whether or not this genus responded to N-addition in a similar manner to ASV-E. Unlike ASV-E, the relative abundance of the entire *Rhizobium* genus did not decrease in response to nitrogen in source soil, bulk soil, or rhizosphere and actually increased in N-addition nodule samples (Fig. 5A). Similarly, N-addition had little effect on Shannon diversity of rhizobia (Fig. 5B), though, similarly to the whole microbial community, diversity decreased strongly across plant compartments: the Shannon diversity of nodules was 15-fold lower (Shannon diversity ∼0.1) than that of the source soil (Shannon diversity ∼1.5) (Fig. 5B, Table S4B), with evenness and richness showing similar patterns (Fig. 5C,D).

**Figure 5:**
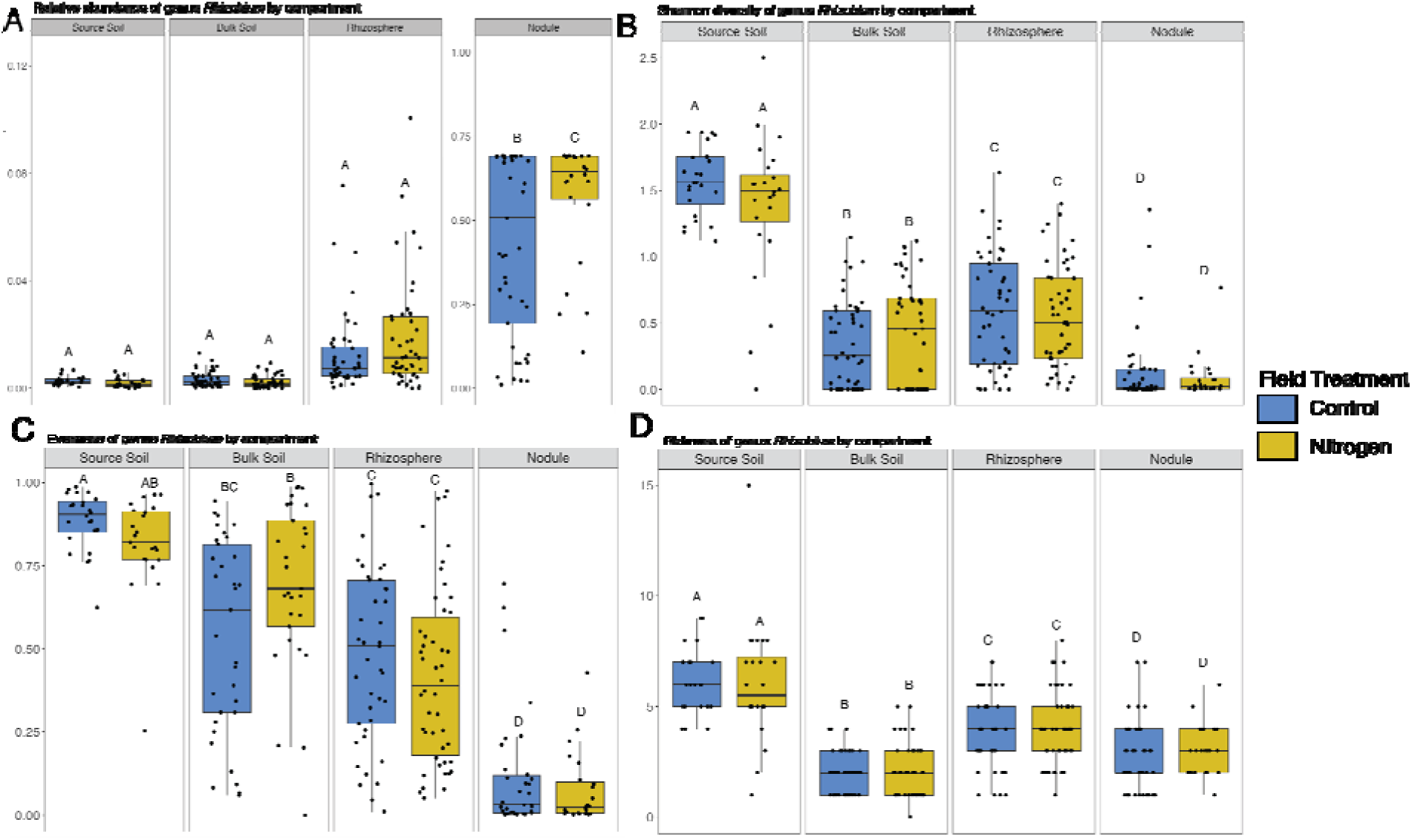
Rhizobium genus relative abundance increases (A) while Rhizobium species diversity and evenness decreases (B,C) within plants compared to soil environments. These shifts in Rhizobium diversity from soil to plant compartments are largely due to increased likelihood of low abundance Rhizobium taxa within rhizosphere (D). A: Log relative abundance of genus *Rhizobium* increases within plants. *P* < 0.05. Significance determined by linear mixed-model and Tukey’s post-hoc test. One upper outlier in the bulk soil (relative abundance 0.29) was removed. Though it was not a statistical outlier, its presence obscured abundance patterns. Letters represent Tukey’s HSD on all 8 Nitrogen x Compartment means (28 comparisons). **B:** Shannon diversity of genus *Rhizobium* significantly decreases within plants. Significance determined by linear mixed-model and Tukey’s post-hoc test. **C**: Evenness of genus *Rhizobium* significantly decreases within plants. *P* < 0.05. Significance determined by linear mixed-model and Tukey’s post-hoc test. Letters represent Tukey’s HSD on all 8 Nitrogen x Compartment means (28 comparisons). 55 values of evenness values were NA, this was because the richness of corresponding samples was 1 and so evenness could not be computed. **D**: Low abundance *Rhizobium* significantly increase richness in source soil, *P* < 0.05. Significance determined by linear mixed-model and Tukey’s post-hoc test.

The relative abundance of *Rhizobium* did not change in response to N, but we did observe significant shifts in community composition within this genus in bulk soil, greenhouse soil, and rhizosphere compartments (nodules were marginal; *P* = 0.07, Table S5, Table S6, Fig. S5, Fig. S6). In all compartments, most ASVs were shared between N-addition and control (Fig. 6A), indicating that ASVs were largely generalists rather than being exclusive to one environment. Nevertheless 21% of all rhizobial ASVs changed in relative abundance between N-addition and control; some increased, while others decreased (Fig. 6B, SD9-10). Four ASVs with the largest increases under N-addition include ASV-B (one of our top nodule ASVs, discussed above), which is significantly more abundant in N-addition in all compartments (Fig. 6B,C). Second, ASV-U (465f73b2a423c62bd060565910acbc79, right next to *R. rhizogenes*; Fig. 4), was never found in the nodules but was significantly more abundant in the three other compartments (Fig. 6B,C). Finally two more ASVs were rare and found only in the source soil (ASV-T: 97f93469eabe0ba1d6933d08c02f1e9d and ASV-M: 9e84fdaea54f2ca70cf83a3afb04af2a; Fig. 6B,C, SD9).

**Figure 6:**
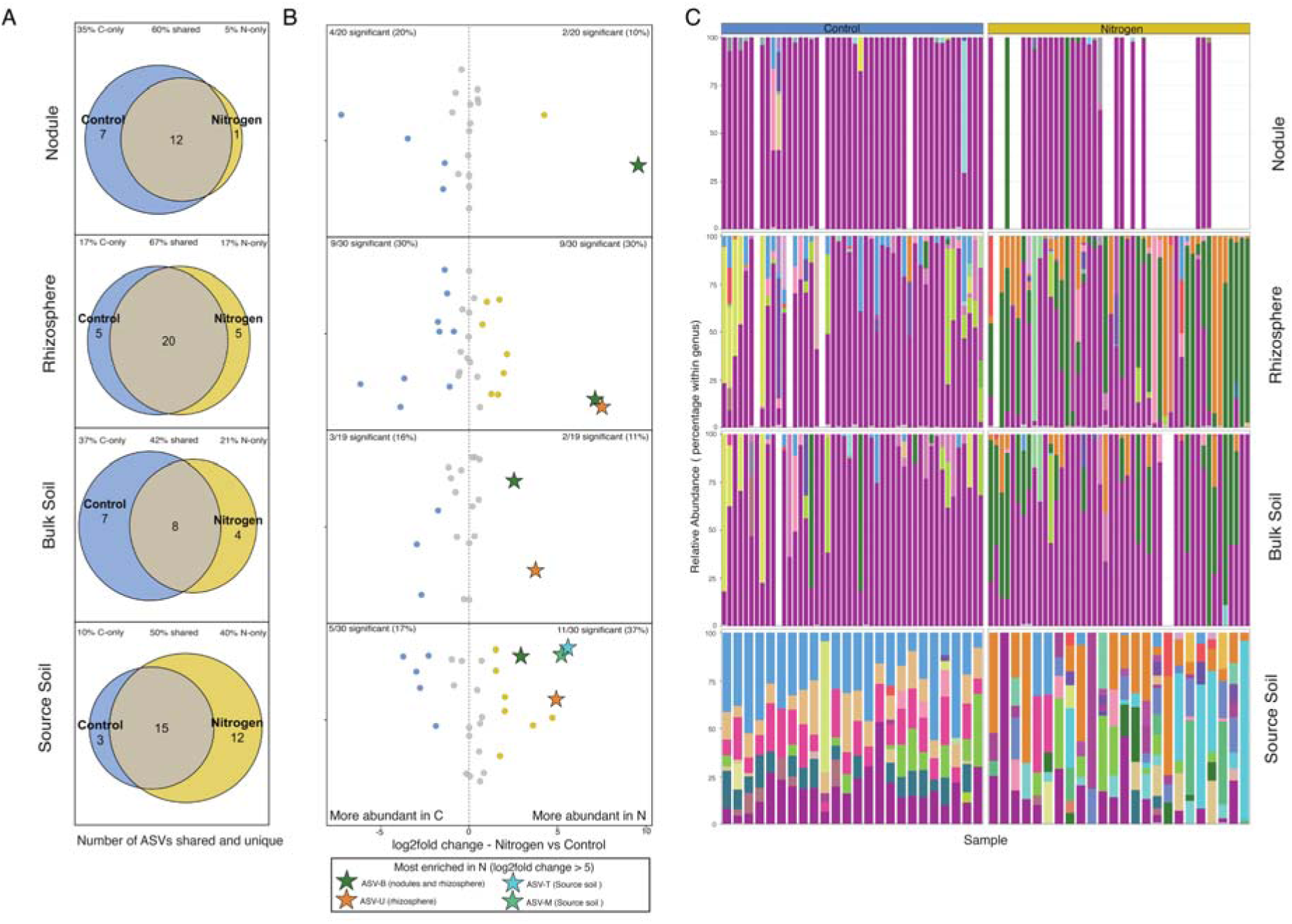
The majority of *Rhizobium* ASVs across compartments are shared between N-addition and control (A), despite strong changes in relative abundance (B,C) A: The majority of *Rhizobium* ASVs are shared between N and. **C**. Compartment-level Venn diagrams depicting whether *Rhizobium* ASVs are unique to N, to C, or are shared between both environments. **B**: Individual *Rhizobium* ASVs respond strongly to N-fertilization within compartments. Log2fold change calculated for 44 *Rhizobium* ASVs across four compartments between nitrogen and control environments. Colored circles represent significant ASVs (p < 0.05), grey circles represent insignificant ASVs (p > 0.05). Four ASVs with log2fold changes greater than 5 in at least one compartment are highlighted with stars instead of dots in every compartment they are present in. C: Species-level diversity of *Rhizobium* is high but varies strongly between N-addition and control. Stacked bar plots of *Rhizobium* ASVs organized by plant compartment. Colors represent unique ASVs; color assignments are consistent within panels. Full ASV identities can be found in Supplementary Data 3-5. ASV relative abundance is normalized within genus. Bars arranged left to right by sample, with the same plant samples being directly above and below each other, all the way to soil core. Samples that are missing, either due to low DNA concentrations or lack of root/nodule are empty white spaces. ASVs that comprise less than 1% of relative abundance in each sample are grouped together and colored grey at the bottom of each box.

## Discussion

Understanding the decline of partner quality in microbial mutualists requires integrative approaches that combine the evolutionary conclusions that can be made from culture-based inoculation experiments with the ecological context provided by community sequencing approaches. Here we reveal that both ecological and evolutionary processes jointly contribute to the breakdown of the agriculturally vital legume-rhizobium mutualism. After 33 years of N-fertilization, a key clover mutualist, *R. leguminosarum* ASV-E, was not only more likely to contain a poor-quality pSym (evolutionary change) but that rhizobial communities shifted such that ASV-E significantly decreased in abundance under long-term N-fertilization (ecological change). Together the outcomes of these processes limit the growth of clover inoculated with microbial communities from long-term N fertilized plots. To add to the ecological complexity of this system, ASV-E was only one member of a diverse rhizobial community in soil and may be outcompeted by these other rhizobia under N-addition. We discuss these major results below.

### Integrative culture-dependent and culture-independent approaches show that ecology and evolution contribute to mutualism decline

In previous work on the breakdown of the legume-*Rhizobium* mutualism, we used a relatively small collection of ∼60 *R. leguminosarum* genospecies E (gsE) and gsB isolates, all isolated from clover nodules, to conclude that N-addition had resulted in evolutionary changes within the rhizobial pSyms resulting in poor partner quality for host plants^38,40^. Here, by inoculating clover plants with soil slurries taken from N-evolved plots 15 years after our initial sampling and leveraging full-length amplicon sequencing of chromosomal and plasmid ASVs, we are able to go beyond this previous culture-dependent work to tease apart the concurrent ecological and evolutionary changes underlying mutualism decline. Not only could we differentiate between closely related *Rhizobium* ASVs at the chromosomal level, unpacking genus-level effects to resolve the community shifts beyond our focal ASVs, we were able to independently demonstrate that the shifts in frequency among the three known pSym clades persist in nature long after our initial sampling.

This convergence across methods and timepoints both validates that our earlier culture-dependent work captured ecologically meaningful variation, while newly demonstrating that these pSym variants have persisted over a timescale during which soil communities will have turned over many times. First, our replication allows us to speak to a recurring tension in microbial ecology over whether culture independent and dependent approaches reach similar conclusions^54,55^, suggesting that, at least for nodule symbionts, the answer is yes. Second, multi-decade monitoring programs have been instrumental in documenting evolutionary change and confirming its persistence in macro-organisms, e.g., Trinidadian guppies^56,57^, Darwin’s finches^58^, but such long-term resampling of microbial communities from nature is exceedingly rare, with long-term work more commonly conducted in laboratories^59^. A few salient examples from nature exist, such as the repeated resampling of stable transconjugants from *Mesorhizobium* associated with *Lotus corniculatus*^60,61^ and the aquatic microbial sampling over two decades in a northern temperate lake^23^. Here we add to that body of literature, concluding that the persistent shifts in the frequencies of clover-associated *Rhizobium* pSyms represent stable evolutionary outcomes rather than transient community fluctuations, with N enrichment as a persistent selective force reshaping the symbiotic qualities of rhizobia.

Our approach allowed us to demonstrate that the biomass effects of ecological changes in N-addition soils (decline in ASV-E due to shifts in the soil *Rhizobium* communities and resulting lack of nodulation) were of similar magnitude to the evolutionary changes (rise in frequency of low-quality pSyms). This suggests that, within the legume-rhizobia mutualism, community-scale access to partners is just as important as within-species partner quality. While past empirical work has shown the importance of symbiont abundance or diversity on host health^62,63^, few studies, if any, compare the magnitude of these effects to within-species evolution of symbiont partner quality. Our work also has interesting implications when considering ways to reverse mutualism breakdown, suggesting that simply replenishing the reservoir of mutualists (even low-quality ones) may improve plant growth. Yet, if the declines in ASV-E result from it being outcompeted by other rhizobial ASVs in soil (ASV-E declines were concomitant with increases in the abundance of other rhizobial ASVs-many of which were not abundant within clover nodules), then supplementing ASV-E into soil may not be effective if the soil community has been restructured in ways that would continue to competitively exclude re-introduced ASV-E^64^. Indeed, because our common garden design captures a snapshot of these communities after 33 years of N-fertilization, we document the outcome of ecological and evolutionary change rather than the field processes that produced it. Multiple mechanisms could have contributed to the clover declines that have been measured in the field^53^ and the patterns we observe here, including reduced plant sanctioning of poor symbionts^11,65^, competitive exclusion of clover by non-legumes that are better at using free N^52^, or the competitive rhizobial dynamics suggested above, and disentangling their relative contributions remains an important direction for future work.

Though resolving ecological and evolutionary processes in microbial communities is vital to understand, since both can separately and jointly change ecosystems and their functioning throughout time^66,67^, our work highlights how this endeavor is still limited by methodological and taxonomic constraints^22,23^. For example, the increased read length and high sample coverage provided by PacBio full-length 16S rRNA sequencing allowed us to make ecological conclusions about mutualism breakdown (ASV-E decreasing in abundance) at the single-ASV level, since the *Rhizobium* genus did not respond in the same way to N-addition as ASV-E. Thus short-read amplicon sequencing, resolving differences at a broader taxonomic level, would have missed this finer-scale change^27,68^. Even so, our methods leave open questions about within-species evolutionary dynamics. We called our top nodule sequence ASV-E due to the fact that it was a perfect full-length match to *R. leguminosarum* genospecies E, one of only two genospecies we have previously cultured from KBS^40^. However, as the 16S rRNA of gsE is identical to the 16S rRNA of ∼10 other genospecies^43^, we are unable to confidently conclude that that this ASV truly represents one taxon. Thus, though we have demonstrated larger-scale ecological change here by showing ASV-E decreasing relative to other diverse *Rhizobium* ASVs, there may be additional evolutionary change occurring *within* the ASV itself (different alleles of gsE or other closely related taxa increasing or decreasing under selection), which we were unable to resolve using our current methods. By examining pSym allele frequencies using a previously identified gsE marker gene, we revealed evolutionary change consistent with selection on gsE pSyms over time. Confirming that these pSym variants are specifically carried by ASV-E, rather than distributed throughout the *Rhizobium* community, requires chromosome-plasmid linkage data from HiC metagenomic sequencing. These difficulties are not unique to our system; in a 20-year freshwater metagenome time series, within-species evolutionary dynamics were also challenging to differentiate from between-species ecological dynamics, which the authors attributed both to technological limitations of sequencing and unclear taxonomic delineations^25^.

### Full-length 16S sequencing resolves rhizobial diversity beyond the limits of database taxonomy

Microbial taxonomy is constantly in flux: reshaped by additional sampling^64,78–80^, inconsistent nomenclature^69,70^ and variable outputs between different analytical tools and pipelines^61,81,8271^. Here, even though we restricted ASVs to the combined genus *Allo/Neo/Para/Rhizobium* (ANPR) within the family *Rhizobiacae* in SILVA, we recovered ASVs spanning across the family, including ASVs which were part of different, SILVA-defined genera than the ANPR genus (such as *Shinella*). The magnitude of this issue was substantial: only 70% of our ASVs were within one of the four ANPR genera, with only 37% of those *Rhizobium.* This likely reflects the fact that many species formerly known as *Rhizobium* sp. are yet to be renamed in public databases such as SILVA even after being reclassified^42,44,69,71^. Despite this, our family-level phylogeny clearly resolved *Rhizobium* sensu lato distinctly from other genera in *Rhizobiaceae,* likely due to both increasing numbers of high-quality references, as well as a significant diversity of ASVs which we could identify due to our increased sequencing resolution. Our data demonstrate that, even as genera themselves continue to be revised, full-length 16S rRNA contains the resolution for genus-level inference when paired with current reference data. As such, we believe that future researchers using full-length 16S rRNA sequencing may benefit from classifying ASVs phylogenetically against curated references, rather than relying solely on large-scale database classifiers, to gain the most possible resolution from the data^72,73^.

Our data also highlight how inconsistent taxonomic labels may obscure the underlying biological processes driving community change. For example, *Rhizobium* is a genus that was originally defined by its function: N-fixing, legume-associated, root nodule bacteria^74^; yet, in our system, plant compartment explained the largest proportion of ANPR ASV variation, with 73% of our 172 ASVs never detected in nodules at all. Also, many of our ASVs, though changing in abundance throughout compartments and under N-addition, were broadly present throughout all compartments and treatments. Though multiple studies have shown rhizobia upregulated in the rhizospheres of non-legumes^75–77^, as well as soil^78^, these data have typically resolved rhizobial taxa at family or genus level, leaving open questions about the presence or abundance of specific symbiont clades, such as clover nodulating *Rhizobium*.

Though our secondary classification has helped resolve taxonomic identity, the majority of the references we used were nodule isolates, while our ASVs were primarily found in the soil and rhizosphere. Our data support the idea that the rhizobia are a flexible, plant-associated group– with nodulation and N-fixation representing just one important life history strategy among potentially many other strategies, such as general plant growth promotion^79^ through secretion of compounds like lipo-chitooligosaccharides (Nod factors)^80^, or potential denitrification (shown in *Bradyrhizobium*^81^). Nevertheless, it is clear that the underlying chromosomal and plasmid diversity of rhizobia remains under-sampled. Additional work integrating metagenomics, long-read amplicon sequencing, and transcriptomics will be key to understanding the distribution and possible functions of rhizobia across plant taxa and soil environments.

## Acknowledgements

We want to acknowledge the technical support of the Roy J Carver Biotechnology Center, particularly Mark Band and Alvaro Hernandez. We also gratefully acknowledge George DiCenzo for providing an up-to-date phylogeny of the family *Rhizobiaceae,* as well as reference sequences, for identification of *Rhizobiaceae* ASVs. Finally, we are extremely thankful for our many collaborators who have contributed useful thoughts to this manuscript in all its forms, including but not limited to David Vereau Gorbitz, Hunter Cobbley, Cari Vanderpool, Caroline Oldstone-Jackson, Ivan Sosa Marquez, Laura Suttenfield, Isaiah Goertz, Jiayue Yang, Sierra Raglin, Tony Yannarell, and Gary Olsen.

## Author contributions

SLB, IML, JAL, RJW, and KDH designed research. SLB, IML, ARH, and AEC conducted research. SLB, KDR, and CJF designed and contributed analytic tools. SLB, IML, CJF, JAL, RJW, KDH analyzed data, and SLB, IML, ARH, CJF, JAL, RJW, and KDH wrote the paper.

## References

1. Boetius, A. et al. A marine microbial consortium apparently mediating anaerobic oxidation of methane. Nature 407, 623–626 (2000).

2. van der Heijden, M. G. A., Bardgett, R. D. & van Straalen, N. M. The unseen majority: soil microbes as drivers of plant diversity and productivity in terrestrial ecosystems. Ecol. Lett. 11, 296–310 (2008).

3. Denison, R. F. Legume Sanctions and the Evolution of Symbiotic Cooperation by Rhizobia. Am. Nat. 156, 567–576 (2000).

4. Beringer, J. E., Brewin, N., Johnston, A. W. B., Schulman, H. M. & Hopwood, D. A. The Rhizobium-Legume Symbiosis. Proc. R. Soc. Lond. B Biol. Sci. 204, 219–233 (1979).

5. Singh, L. P., Gill, S. S. & Tuteja, N. Unraveling the role of fungal symbionts in plant abiotic stress tolerance. Plant Signal. Behav. 6, 175–191 (2011).

6. Márquez, L. M., Redman, R. S., Rodriguez, R. J. & Roossinck, M. J. A Virus in a Fungus in a Plant: Three-Way Symbiosis Required for Thermal Tolerance. Science 315, 513–515 (2007).

7. Scott, J. J. et al. Bacterial Protection of Beetle-Fungus Mutualism. Science 322, 63–63 (2008).

8. Boucher, D. H., James, S. & Keeler, K. H. The Ecology of Mutualism. Annu. Rev. Ecol. Syst. 13, 315–347 (1982).

9. Bronstein, J. L. Our Current Understanding of Mutualism. Q. Rev. Biol. 69, 31–51 (1994).

10. Dethlefsen, L., McFall-Ngai, M. & Relman, D. A. An ecological and evolutionary perspective on human–microbe mutualism and disease. Nature 449, 811–818 (2007).

11. Sachs, J. L. & Simms, E. L. Pathways to mutualism breakdown. Trends Ecol. Evol. 21, 585–592 (2006).

12. Kiers, E.T., Palmer, T. M., Ives, A. R., Bruno, J. F. & Bronstein, J. L. Mutualisms in a changing world: an evolutionary perspective. Ecol. Lett. 13, 1459–1474 (2010).

13. Briand, F. & Yodzis, P. The Phylogenetic Distribution of Obligate Mutualism: Evidence of Limiting Similarity and Global Instability. Oikos 39, 273–275 (1982).

14. Gano-Cohen, K. A. et al. Recurrent mutualism breakdown events in a legume rhizobia metapopulation. Proc. R. Soc. B Biol. Sci. 287, (2020).

15. Hays, B. R. et al. Demographic consequences of mutualism disruption: Browsing and big-headed ant invasion drive acacia population declines. Ecology 103, e3655 (2022).

16. Palmer, T. M. et al. Breakdown of an Ant-Plant Mutualism Follows the Loss of Large Herbivores from an African Savanna. Science 319, 192–195 (2008).

17. Bay, R. A. & Palumbi, S. R. Multilocus Adaptation Associated with Heat Resistance in Reef-Building Corals. Curr. Biol. 24, 2952–2956 (2014).

18. Selmoni, O., Cleves, P. A. & Exposito-Alonso, M. Global coral genomic vulnerability explains recent reef losses. Nat. Commun. 17, 896 (2025).

19. Zhang, T. et al. The evolution of parasitism from mutualism in wasps pollinating the fig, Ficus microcarpa, in Yunnan Province, China. Proc. Natl. Acad. Sci. 118, e2021148118 (2021).

20. Jones, E. I., Ferrière, R. & Bronstein, J. L. Eco-evolutionary dynamics of mutualists and exploiters. Am. Nat. 174, 780–794 (2009).

21. Weinbach, A., Loeuille, N. & Rohr, R. P. Eco-evolutionary dynamics further weakens mutualistic interaction and coexistence under population decline. Evol. Ecol. 36, 373–387 (2022).

22. Batarseh, T. N. & Koskella, B. Distinguishing among evolutionary and ecological processes shaping microbiome dynamics. ISME J. 19, wraf107 (2025).

23. Rohwer, R. R. et al. Two decades of bacterial ecology and evolution in a freshwater lake. Nat. Microbiol. 10, 246–257 (2025).

24. Lewis, W. H., Tahon, G., Geesink, P., Sousa, D. Z. & Ettema, T. J. G. Innovations to culturing the uncultured microbial majority. Nat. Rev. Microbiol. 19, 225–240 (2021).

25. Cordero, O. X. & Polz, M. F. Explaining microbial genomic diversity in light of evolutionary ecology. Nat. Rev. Microbiol. 12, 263–273 (2014).

26. VanInsberghe, D., Arevalo, P., Chien, D. & Polz, M. F. How can microbial population genomics inform community ecology? Philos. Trans. R. Soc. B Biol. Sci. 375, 20190253 (2020).

27. Johnson, J. S. et al. Evaluation of 16S rRNA gene sequencing for species and strain-level microbiome analysis. Nat. Commun. 10, 5029 (2019).

28. Tedersoo, L., Albertsen, M., Anslan, S. & Callahan, B. Perspectives and Benefits of High-Throughput Long-Read Sequencing in Microbial Ecology. Appl. Environ. Microbiol. 87, e00626–21.

29. Weisberg, A. J. et al. Unexpected conservation and global transmission of agrobacterial virulence plasmids. Science 368, eaba5256 (2020).

30. Polz, M. F., Alm, E. J. & Hanage, W. P. Horizontal Gene Transfer and the Evolution of Bacterial and Archaeal Population Structure. Trends Genet. TIG 29, 170–175 (2013).

31. Wardell, G. E., Hynes, M. F., Young, P. J. & Harrison, E. Why are rhizobial symbiosis genes mobile? Philos. Trans. R. Soc. B Biol. Sci. 377, 20200471 (2022).

32. Heath, K. D., Batstone, R. T., Cerón Romero, M. & McMullen, J. G. MGEs as the MVPs of Partner Quality Variation in Legume-Rhizobium Symbiosis. mBio 13, e00888–22 (2022).

33. López-Madrigal, S. & Gil, R. Et tu, Brute? Not Even Intracellular Mutualistic Symbionts Escape Horizontal Gene Transfer. Genes 8, 247 (2017).

34. Wilkinson, D. M. & Sherratt, T. N. Horizontally acquired mutualisms, an unsolved problem in ecology? Oikos 92, 377–384 (2001).

35. Drew, G. C., Stevens, E. J. & King, K. C. Microbial evolution and transitions along the parasite–mutualist continuum. Nat. Rev. Microbiol. 19, 623–638 (2021).

36. Martiny, J. B. H. et al. Investigating the eco-evolutionary response of microbiomes to environmental change. Ecol. Lett. 26, S81–S90 (2023).

37. Turgeon, B. G. & Bauer, W. D. Ultrastructure of infection-thread development during the infection of soybean by Rhizobium japonicum. Planta 163, 328–349 (1985).

38. Weese, D. J., Heath, K. D., Dentinger, B. T. M. & Lau, J. A. Long-term nitrogen addition causes the evolution of less-cooperative mutualists. Evolution 69, 631–642 (2015).

39. Klinger, C. R., Lau, J. A. & Heath, K. D. Ecological genomics of mutualism decline in nitrogen-fixing bacteria. Proc. R. Soc. B Biol. Sci. 283, 20152563 (2016).

40. Vereau Gorbitz, D., et al. Plasmid transmission dynamics and evolution of partner quality in a natural population of Rhizobium leguminosarum. mBio 0, e02497–25 (2025).

41. Sawada, H., Kuykendall, L. D. & Young, J. M. Changing concepts in the systematics of bacterial nitrogen-fixing legume symbionts. J. Gen. Appl. Microbiol. 49, 155–179 (2003).

42. Naranjo-Robayo, N. et al. Updated taxonomy of the family Rhizobiaceae with proposals for 10 novel genera and 35 novel combinations. 2025.11.20.689363 Preprint at 10.1101/2025.11.20.689363 (2025).

43. Young, J. P. W. et al. Defining the Rhizobium leguminosarum Species Complex. Genes 12, 111 (2021).

44. Kuzmanović, N., Fagorzi, C., Mengoni, A., Lassalle, F. & diCenzo, G. C. Taxonomy of Rhizobiaceae revisited: proposal of a new framework for genus delimitation. Int. J. Syst. Evol. Microbiol. 72, 005243 (2022).

45. Weisberg, A. J., Miller, M., Ream, W., Grünwald, N. J. & Chang, J. H. Diversification of plasmids in a genus of pathogenic and nitrogen-fixing bacteria. Philos. Trans. R. Soc. B Biol. Sci. 377, 20200466 (2021).

46. Hollowell, A. C. et al. Epidemic Spread of Symbiotic and Non-Symbiotic Bradyrhizobium Genotypes Across California. Microb. Ecol. 71, 700–710 (2016).

47. Sachs, J. L., Kembel, S. W., Lau, A. H. & Simms, E. L. In Situ Phylogenetic Structure and Diversity of Wild Bradyrhizobium Communities. Appl. Environ. Microbiol. 75, 4727–4735 (2009).

48. Laguerre, G., Bardin, M. & Amarger, N. Isolation from soil of symbiotic and nonsymbiotic *Rhizobium leguminosarum* by DNA hybridization. Can. J. Microbiol. 39, 1142–1149 (1993).

49. Zézé, A., Mutch, L. A. & Young, J. P. W. Direct amplification of nodD from community DNA reveals the genetic diversity of Rhizobium leguminosarum in soil. Environ. Microbiol. 3, 363–370 (2001).

50. Sarita, S., Sharma, P. K., Priefer, U. B. & Prell, J. Direct amplification of rhizobial nodC sequences from soil total DNA and comparison to nodC diversity of root nodule isolates. FEMS Microbiol. Ecol. 54, 1–11 (2005).

51. Huberty, L. E., Gross, K. L. & Miller, C. J. Effects of nitrogen addition on successional dynamics and species diversity in Michigan old-fields. J. Ecol. 86, 794–803 (1998).

52. Dickson, T. L. & Gross, K. L. Plant community responses to long-term fertilization: changes in functional group abundance drive changes in species richness. Oecologia 173, 1513–1520 (2013).

53. Kay Gross & Jennifer Lau. Plant Community and Ecosystem Responses to Long-term Fertilization and Disturbance at the Kellogg Biological Station. Environmental Data Initiative 10.6073/pasta/ea22c735ddfe17c595ccca978a87d109.

54. Martiny, A. C. High proportions of bacteria are culturable across major biomes. ISME J. 13, 2125–2128 (2019).

55. Steen, A. D. et al. High proportions of bacteria and archaea across most biomes remain uncultured. ISME J. 13, 3126–3130 (2019).

56. Reznick, D. N., Shaw, F. H., Rodd, F. H. & Shaw, R. G. Evaluation of the Rate of Evolution in Natural Populations of Guppies (Poecilia reticulata). Science 275, 1934–1937 (1997).

57. Reznick, D. N. & Ghalambor, C. K. Selection in Nature: Experimental Manipulations of Natural Populations. Integr. Comp. Biol. 45, 456–462 (2005).

58. Grant, P. R. & Grant, B. R. Unpredictable evolution in a 30-year study of Darwin’s finches. Science 296, 707–711 (2002).

59. Lenski, R. E. Experimental evolution and the dynamics of adaptation and genome evolution in microbial populations. ISME J. 11, 2181–2194 (2017).

60. Sullivan, J. T., Patrick, H. N., Lowther, W. L., Scott, D. B. & Ronson, C. W. Nodulating strains of Rhizobium loti arise through chromosomal symbiotic gene transfer in the environment. Proc. Natl. Acad. Sci. 92, 8985–8989 (1995).

61. Colombi, E. et al. Population genomics of Australian indigenous Mesorhizobium reveals diverse nonsymbiotic genospecies capable of nitrogen-fixing symbioses following horizontal gene transfer. *Microb*. Genomics 9, mgen000918 (2023).

62. Burghardt, L. T., Epstein, B., Hoge, M., Trujillo, D. I. & Tiffin, P. Host-Associated Rhizobial Fitness: Dependence on Nitrogen, Density, Community Complexity, and Legume Genotype. Appl. Environ. Microbiol. 88, e00526–22 (2022).

63. Dunkley, K., Cable, J. & Perkins, S. E. Consistency in mutualism relies on local, rather than wider community biodiversity. Sci. Rep. 10, 1–11 (2020).

64. Mendoza-Suárez, M., Andersen, S. U., Poole, P. S. & Sánchez-Cañizares, C. Competition, Nodule Occupancy, and Persistence of Inoculant Strains: Key Factors in the Rhizobium-Legume Symbioses. Front. Plant Sci. 12, 690567 (2021).

65. Kiers, E. T., Rousseau, R. A., West, S. A. & Denison, R. F. Host sanctions and the legume–rhizobium mutualism. Nature 425, 78–81 (2003).

66. Walsh, M. R., DeLong, J. P., Hanley, T. C. & Post, D. M. A cascade of evolutionary change alters consumer-resource dynamics and ecosystem function. Proc. R. Soc. B Biol. Sci. 279, 3184–3192 (2012).

67. Post, D. M. & Palkovacs, E. P. Eco-evolutionary feedbacks in community and ecosystem ecology: interactions between the ecological theatre and the evolutionary play. Philos. Trans. R. Soc. B Biol. Sci. 364, 1629–1640 (2009).

68. Yarza, P. et al. Uniting the classification of cultured and uncultured bacteria and archaea using 16S rRNA gene sequences. Nat. Rev. Microbiol. 12, 635–645 (2014).

69. Lloyd, K. G. & Tahon, G. Science depends on nomenclature, but nomenclature is not science. Nat. Rev. Microbiol. 20, 123–124 (2022).

70. Bobay, L.-M. The Prokaryotic Species Concept and Challenges. in The Pangenome: Diversity, Dynamics and Evolution of Genomes (eds Tettelin, H. & Medini, D.) (Springer, Cham (CH), 2020).

71. Edgar, R. Taxonomy annotation and guide tree errors in 16S rRNA databases. PeerJ 6, e5030 (2018).

72. Dueholm, M. S. et al. Generation of Comprehensive Ecosystem-Specific Reference Databases with Species-Level Resolution by High-Throughput Full-Length 16S rRNA Gene Sequencing and Automated Taxonomy Assignment (AutoTax). mBio 11, e01557–20 (2020).

73. Myer, P. R. et al. Classification of 16S rRNA reads is improved using a niche-specific database constructed by near-full length sequencing. PLOS ONE 15, e0235498 (2020).

74. Root Nodule Bacteria and Leguminous Plants. https://www.library.wisc.edu/parallelpress/pp-catalog/books/root-nodule-bacteria-and-leguminous-plants/.

75. Kumar, M. et al. Core microbiota of wheat rhizosphere under Upper Indo-Gangetic plains and their response to soil physicochemical properties. Front. Plant Sci. 14, 1186162 (2023).

76. Yeoh, Y. K. et al. Evolutionary conservation of a core root microbiome across plant phyla along a tropical soil chronosequence. Nat. Commun. 8, 215 (2017).

77. Xu, J. et al. The structure and function of the global citrus rhizosphere microbiome. Nat. Commun. 9, 4894 (2018).

78. Jones, F. P. et al. Novel European free-living, non-diazotrophic Bradyrhizobium isolates from contrasting soils that lack nodulation and nitrogen fixation genes – a genome comparison. Sci. Rep. 6, 25858 (2016).

79. Fahde, S., Boughribil, S., Sijilmassi, B. & Amri, A. Rhizobia: A Promising Source of Plant Growth-Promoting Molecules and Their Non-Legume Interactions: Examining Applications and Mechanisms. Agriculture 13, 1279 (2023).

80. Tanaka, K. et al. Effect of lipo-chitooligosaccharide on early growth of C4 grass seedlings. J. Exp. Bot. 66, 5727–5738 (2015).

81. Woliy, K., Degefu, T. & Frostegård, Å. Host Range and Symbiotic Effectiveness of N2O Reducing Bradyrhizobium Strains. Front. Microbiol. 10, (2019).

