## Supplementary information for "Concurrent ecological and evolutionary processes contribute to mutualism breakdown between legumes and rhizobia"

**Supplementary Methods**

**Long-term ecological site and soil collection details:** The KBS plots are fertilized yearly in June with 12.3 g N/m^-2^ annually in the form of ammonium nitrate or urea. All three *Trifolium* species, *T. repens, T. hybridum,* and *T. pratense*, are frequently found in the control plots but are almost absent from the N-fertilized plots^51-53^. In the six replicate plots, for both the Nitrogen addition and unfertilized treatment subplots, four soil cores (10 mm diameter, 5 cm deep) were collected after removing the top 1 cm to reduce plant debris. Soils were homogenized after removal of the top centimeter and stored at 4°C until inoculation. Cores were taken 1.67 m in from both the x and y axes (equivalent to 2.36 m diagonally from each corner) to ensure even spatial distribution and to avoid edge effects.

**Greenhouse growth details:** Seeds were planted on July 20, 2022, inoculated on July 27, 2022, and harvested the week of September 19, 2022 (at an actively growing stage prior to bud or flower formation). Plants were grown in the University of Illinois Greenhouse (approximate coordinates 40.099155°N, 88.223847°W). Plants were grown in sterile conetainers (Ray Leach Super Cell Classic SC10, 1.5 x 8.25, Steuwe and Sons, Oregon USA) containing a sterile substrate mixture of ½ turface, ¼ coarse quartz sand and ¼ torpedo sand, and topped with vermiculite to retain moisture. During the experiment, plants received natural light (approximately 12-14 hours/day), and water by misting (twice a day, 15 minutes each). Clover seeds were sourced from Ernst Seeds, PA, USA.

Soil slurry inocula were made by combining 5g of field soil (collected 24 hours prior and kept at 4°C) with 45 of 1X PBS; 2 ml of the appropriate inoculum was inoculated onto each plant following previous protocols^38^. Plants were also provided with 10 mL of N-free Fahraeus micronutrient solution weekly and no other nutrients^S1^. Plants were spaced out on greenhouse trays to minimize leaf contact.

Each of the three identical blocks contained two plants (one of each host species) inoculated with each soil core, as well as 30 controls: 16 pots inoculated with autoclaved soil, 10 with sterile PBS, and 4 soil-only “no-plant” pots. This resulted in 126 pots per block and 398 pots in total ({48 soil inocula x 2 host plant species + 30 controls per block} x 3 blocks). Given this design, replication of the microbial community effect comes from having multiple cores per plot, and multiple plots per treatment, rather than within-core replication, allowing us to maximize observable variation and maintain tractability.

Virtually no contamination was observed; control plants rarely produced nodules and always had biomass below 5mg (Supplementary Data 1). For our plants inoculated with autoclaved soil (to control for abiotic effects between N-addition and control soil at KBS, we observed extremely low biomass values, with no differences between plants inoculated with N-addition and control soils, suggesting that the biomass differences of our plants inoculated with live soil cannot be explained by residual soil N or other minerals (Fig. S7).

**Harvest and processing details:** Plants were harvested after eight weeks of growth at a vegetative growth stage prior to flowering. Aboveground biomass was taken after drying for ~7 days at 65°C. For bulk soil, we split the Falcon tube into two cryotubes and froze them at -80°C. We also froze the remainder of the Falcon tube for other possible analyses. Rhizosphere samples were collected using a modified published protocol^S2^; we added 100uL of Tween20 (Sigma-Aldrich, MO, USA) to each root tube, incubated at room temperature for 20 minutes, and then vortexed for 1 minute at 14,000 RPM. Then we removed roots, pelleted the mixture, and froze it at -80°C. Finally, we plucked ten random nodules off of each root containing nodules, choosing a mixture of large, pink nodules and smaller, whiter nodules. We processed nodules within 24H after harvesting to minimize the possibility of senescence, as we could not tell the difference between non-functional nodules and previously functional, senesced nodules. We sterilized them by first dipping in 70% ethanol for 30 seconds, rinsing in sterile water, and then dipping in 10% bleach for 30 seconds^S3^. All samples remained frozen until used for DNA extraction.

All three blocks were processed within three days of the initial harvest. Blocks 2 and 3 were frozen at -80°C, and the nodules of Block 1 were counted fresh (within 1 week in total.

**Modifications to DNA extraction protocol:** We extracted DNA from all samples using Qiagen’s PowerSoil Pro Kit (Qiagen, Hilden, Germany) in sterile flow hoods. All DNA extracts were eluted to 100uL. For greenhouse bulk soil, we used 500 mg of soil. For rhizosphere, we used the entire rhizosphere pellet. For nodules, we homogenized ten sampled nodules from each plant containing nodules.

For the greenhouse bulk soil only, we modified Step 1 by combining 300 μL of ATL buffer (Qiagen, Hilden, Germany) and 500uL of CD1, rather than using 800 μL of CD1. This was to assist in the removal of negatively-charged DNA from positively-charged Turface clay, where it had strongly adsorbed. We then continued the protocol as written after Step 1. For nodule samples only, we modified Steps 1-2 by adding 800 μL of CD1 directly to the tube containing the nodules then homogenizing nodules with a pestle.

**Library prep and sequencing details:**

In total, we sequenced 364 16S rRNA samples (48 source soil, 123 bulk soil, 124 rhizosphere, and 69 nodule samples) and 69 *nodA* samples (the same nodule DNA used for 16S rRNA). Nearly every missing sample was from N-plot plants that either died or weighed less than 1mg, so they likely never formed nodules or roots large enough to sequence (Supplementary Data 11-12). The full-length 16S amplicons were generated from 2.5ng of DNA with the barcoded Full-Length (27F-1492R) Kinnex 16S primers from PacBio (forward 5’-AGRGTTYGATYMTGGCTCAG-3’ and reverse 5’-RGYTACCTTGTTACGACTT-3’, producing ~1460 bp amplicons) and the 2x Roche KAPA HiFi Hot Start Ready Mix following the protocol from PacBio. PCR was set up on ice in 25uL volumes using barcoded primers with the following reagents: 12uL Kappa Hotstart Ready Mix (Roche Biosciences, Basel, Switzerland), 1.5 μL water.,3uL forward primer, 3uL reverse primer, 5uL DNA (0.5 ng/uL). Primers have a 5’ barcode and Kinnex linker extension. PCR cycling parameters were: 95°C denaturing, 3min. 25 cycles of 95°C, 30 seconds, 57°C, 30 seconds, 72°C, 60 seconds. Cycle number was restricted to 25 to reduce probability of chimera generation.

The *nodA* amplicons were generated using published primers^S4^ with the forward primer 5’-TGCRGTGGAARNTRNNCTGGGAAA and with the reverse primer 5'-GGNCCGTCRTCRAAWGTCARGTA (amplifying ~650 bp) and amplified with a 2x Roche KAPA HiFi Hot Start Ready Mix enzyme. PCR was set up on ice in 25uL volumes using barcoded primers with the following reagents: 12.5 μL Kappa Hotstart Ready Mix (KAPA Biosystems), 1.5 μL water, 3 μL forward primer, 3 μL reverse primer, 5 μL DNA (0.5 ng/uL). Primers have Fluidigm 5’ CS1 and CS2 linkers. PCR cycling parameters were: 95°C denaturing, 3min. 28 cycles of 95°C, 30 seconds, 57°C, 30 seconds, 72°C, 60 seconds. Cycle number was restricted to 28 to reduce probability of chimera generation. PCR from the first round was then diluted 1:70 with water and 1ul was used as a template for the barcoding reaction. This was set up using 6.2 μL Kappa Hotstart Ready Mix (KAPA Biosystems), 2.8uL water, 2.4 μL Fluidigm barcodes, and 1uL diluted PCR product. PCR cycling parameters were: 95°C denaturing, 3 minutes. 14 cycles of 95°C, 30 seconds, 57°C, 30 seconds, 72°C 60 seconds.

For both 16S rRNA and *nodA,* all products were measured using SpectraMaxQuant AccuClear Nano kit (Molecular Devices, CA, USA) and run on a fragment analyzer for QC. Samples were pooled according to concentration and cleaned twice with 0.6 volumes of magnetic beads. The amplicons were concatenated 12-fold with the PacBio Kinnex Concatenation Kit and Kinnex PCR 12-Fold Kit to produce an 18kb library. The concatenated library was quantitated with Qubit and the run on a Fragment Analyzer (Agilent, CA, USA) to confirm the presence of DNA fragments of the expected size. Sequencing was completed on the PacBio Revio instrument with a 30 hr movie time. Circular Consensus analysis (CCS), de-concatenation, and demultiplexing were done using SMRTlink 13.1 with RQ>=0.999.

**Sequence processing details:** PacBio HiFi reads were trimmed using cutadapt v4.1. For 16S rRNA, the 27F primer is reverse complemented based on cutadapt specifications (specifications can be found on Zenodo; 10.5281/zenodo.17479679). For nodA, trimming utilized cutadapt but using *nodA* primer sequences described above. Trimmed reads were processed using DADA2 v.1.30.0^S5,6^. Potential chimeric sequences were detected and removed using removeBimeraDenovo. To identify *nodA* sequences, aligned sequences of the full *nodA* protein family from the Protein families (Pfam) database hosted on Interpro were downloaded in Stockholm format^S7^. The profile was initially converted to a simple FASTA file prior to formatting for profile searching and finally to a tab-delimited file (BLAST ‘m8’ format). ASV sequences were filtered to only retain those with a highly significant match (e-value of less than 1e-5) to at least one protein from the *nodA* profile search results. Pipeline can be found on Zenodo (10.5281/zenodo.17479679).

For 16S rRNA, taxonomic assignments were performed using the DADA2 implementation of the RDP classifier^S8^ and the SILVA 138.1 reference database^S9,S10^ including species-level assignments. For *nodA,* the above steps were performed with the following changes. Standard taxonomic screening using the DADA2 implementation of the RDP classifier was not performed. Instead, resulting ASV sequences were screened using MMseqs2 v17.b804f^S11^ to identify positive *nodA* sequences by profile-based alignment and remove any potential background amplification artifacts.

The *nodA* primers used were general enough to allow for amplification of less conserved sequences, both from other more distantly related rhizobia and other known N-fixers such as *Cupriavidus*, which we had observed at the 16S level as dominant in multiple nodule samples (Fig. 1A, Fig. S4).

**Initial sequence analysis and filtering:** We recovered ~162,000 ASVs from the initial run, with a mean read count of 158,000 reads/sample (Supplementary Data 13). We removed mitochondrial and chloroplast contaminants by removing all ASVs at the family *Mitochondria* as well as all within the order *Chloroplast.* Because rarefying discards valid reads and inflates variance, we instead applied targeted prevalence and depth filters following current recommendations^S12–14^: 1) ASVs needed to be present in at least two samples. 2) ASVs needed to be present at a depth of at 100 reads or more in at least one sample they occurred in.

Additionally, we observed in our nodule samples a particularly high prevalence of rare, closely related (usually 1 SNP from each other), and highly co-occurring sequences. These ASVs would often appear in large groups, for example, a group of ~30 ASVs all present in between 50-150 reads each in exactly the same 25 samples. Upon viewing of a multiple sequence alignment (MSA) of these sequences, it became apparent that many of these sequences represented an apparent chimera formed from an abundant parent an d containing 1-2 SNPs that were only found in chloroplast/mitochondrial contaminants. We believe these reads represent sequencing error; however, these were not captured by more traditional chimeric removal steps. To address this issue, we developed an algorithm to filter out biological diversity from sequencing error.

This algorithm, available on Zenodo ( 10.5281/zenodo.17479679), took a multiple sequence alignment (MSA) of all nodule ASVs (including rare ASVs, and chloroplast/mitochondrial contaminants), as well as individual files of raw ASV reads and sample presence/absence, and an input database of known 16S rRNA variants in *Rhizobium*. The algorithm uses ASV rarity, abundant parent identity, and SNP identity to determine whether or not a nodule ASV is likely to be biologically real or not. As a useful sanity check, the vast majority of ASVs called “likely artifact” (3909) were also ASVs which we separately filtered out due to low abundance or prevalence (3767). We removed all ASVs classified as “unclear” or “likely artifact”, only maintaining ASVs that were classified as “likely real”. We then merged this data frame with the first two classifiers for rarity and sample presence, to have a final ASV dataset consisting of 1) ASVs present in more than one sample, 2) ASVs present in at least 100 reads in at least one sample, and 3) if the ASV is present in nodules, passing the chimera filter. Together, these criteria removed most artificial sequences while preserving authentic community diversity, including nodule specialists. After filtering we were left with 7759 ASVs belonging to 389 genera (as compared to 7836 ASVs when we performed the same sample number and prevalence filtering but additionally filtered out all ASVs that were only observed in the nodules).

***Rhizobium* classification:**

Using the *Rhizobiaceae* reference sequences, we constructed a phylogenetic tree of the family. First, we roughly aligned all ASVs and references in Geneious, which allowed us to trim all references and ASVs to the same length. We then extracted the individual sequences and aligned again using MAFFT^S15^ on the local pair settings with 1000 iterations on 30 threads. We used IQ-TREE2^S16^ to infer phylogenetic relationships. The aligned sequences were analyzed using 1000 ultrafast bootstrap replicates (-B 1000) to assess branch support. Standard settings were employed; IQ-TREE2 automatically selected the best-fit substitution model. We then visualized and annotated the tree using iTOL. Genus annotations on iTOL were derived from recent genera propositions^42^, so some species names may appear inaccurate (*e.g.,* a reference called *Rhizobium* in a clade that is not *Rhizobium*) if they have been proposed to be reclassified. Nodule presence was defined as an ASV being present in at least one nodule sample. Plant specificity was defined as an ASV only being present in rhizosphere and/or nodule samples, and never in bulk soil and/or source soil. Full genera annotations can be viewed on iTOL: <https://itol.embl.de/tree/1041903595289581779643614>.

To further classify specific ANPR ASVs (those in Fig. 1) below the genus level, we downloaded genomes from the newest *Rhizobiaceae* taxonomic classification^42^ (see Supplementary Data 5 for accessions). We extracted the 16S rRNA copies from each genome using Barrnap^S17^, retaining all copies over 1400 nucleotides as many *Rhizobium* species have non-clonal 16S rRNA. We employed BLAST on standard settings to map ASVs to their best hits and kept all full-length matches. All matches were above 1264 base pairs, and none were discarded due to short length. To further classify these ASVS into genospecies, we downloaded all reference *Rlc* genospecies^43^, extracted the 16S rRNA, and employed BLAST using the same metrics. Matches can be found in Supplementary Data 5. Code can be found on Zenodo (10.5281/zenodo.17410936).

***Data analysis***

*Microbial community responses to field N-fertilization treatments:* To test whether the community composition of 1) the whole microbial community, 2) the *ANPR* genera, and 3) the *Rhizobium* senso latu differed between nitrogen treatment, plant compartment (source soil, bulk soil, rhizosphere, or nodule), plant species, and two-way interactions/three-way between the fixed factors, we used PERMANOVA (vegan, adonis2, by = terms) on Bray-Curtis dissimilarities of relative ASV abundances. Plot (6 plots per field nitrogen treatment) was treated as a stratified factor. Because the initial PERMANOVA indicated a compartment x nitrogen interaction, we also performed compartment-specific PERMANOVAs. Visualization of these dissimilarities was performed using distance-based redundancy analysis (dbRDA) (capscale) using the same model as the PERMANOVA.

To examine the composition of nodules, we created a stacked bar plot of all genera within nodules, as well as a pie chart of all ANPR ASVs in nodules. For the stacked bar plot, ASVs were grouped at the genus level and their relative abundances between samples were calculated, then each sample’s compositions were averaged to make an overall plot of relative abundances. For both figures, genera/ASVs with a mean relative abundance of less than 1% overall were grouped into “Other”. A Fisher’s exact test was performed to test whether the proportion of nodules dominated by ANPR differed between N-addition and control

We tested how nitrogen affected *Rhizobium* relative abundance and genus diversity (species richness, Shannon diversity, and evenness) with a linear mixed-effects model with field nitrogen treatment, compartment, and the interaction of nitrogen and compartment as fixed predictor variables and plot as a random factor(lmerTest^S18^). Because source soil samples were not part of the greenhouse design/had no plant species associated, we computed separate models that removed these samples and included plant species as a fixed effect. Plant species had no effect on relative abundance or evenness. Species main effects on richness and Shannon diversity were consistent with the species-level differences observed in plant biomass, and did not interact with nitrogen treatment; these results are not further reported.

*Differential abundance of Rhizobium ASVs and nodA alleles:* We used DESeq2^S19^ v1.48.1 testing the individual compartment-specific effects of nitrogen treatment (Nitrogen treatment * Compartment) (for *Rhizobium* ASVs) and nitrogen treatment alone (for *nodA* alleles). P-values were adjusted by the Benjamini-Hochberg procedure, and ASVs/alleles with an FDR-corrected p value <0.05 were deemed significant. As our analysis was restricted to a small taxonomic subset (one genus), we used a +1 pseudocount rather than the built-in poscounts method. Because rhizobial relative abundance did not differ significantly between N and C in soil and rhizosphere compartments (Fig. 4A), nor did library size for *nodA,* the asymmetric size factors produced by poscounts can interpret biological abundance differences as differences in technical library size. We validated our approaches against the poscounts method; overall conclusions did not change (results not further reported). The four chromosomal ASVs with fold changes above 5 were starred with the same color as their corresponding ASVs to the right (Fig. 5A and 5C).

*Identifying individual microbial taxa and pSym alleles resulting in mutualism decline (reduced host benefit):* To test whether ASV-E abundance was significantly different under N vs C, we fit four separate linear mixed models with lmer (Log_Abundance ~ Nitrogen_treat+ 1|Plot), although plot did not explain any variation in the nodule compartment. Fixed-effects F-tests were type III with Kenward-Rodger degrees of freedom. As nitrogen was the only fixed term (no compartment factor in the individual models), the Type III test is equivalent to a standard Type I analysis, but was retained to maintain consistency with other models. Models were fitted separately for each plant compartment, as relative abundance within nodules was so much higher that it dwarfed meaningful variation in other compartments. To test whether nodulation was higher when plants were inoculated with control plot soil slurries, a Fisher’s exact test was used on the 2x2 contingency table of nitrogen treatment by nodulation (present/absent). To test whether shoot biomass differed between nodules dominated by pSym type 3 and all other nodules, we used a Wilcoxon rank-sum test.

*Piecewise structural equation model (SEM)*

To test the pathway from soil community composition to plant biomass, we employed a two-part piecewise SEM^S20^ corresponding to the sequential biological effects of nodulation and symbiont quality. First, to test whether ASV-E explained nodulation, we fit a logistic regression on the source soil relative abundance of ASV-E (n = 48) predicting a binary of nodule presence/absence on all soil cores. We transformed the nodule number of all nodulated plants into presence/absence, and then calculated the mean per soil core (so, if any plant inoculated with a core had nodulated, the core would be scored with a 1. If no plants in a core nodulated, the core would be scored a 0). Then, to test the proportion of biomass which was driven by pSym dominance and nodulation, we fit a linear model (n=48) on soil cores with two predictors: a binary indicator on whether the core had nodulated, and a nested binary indicator, scored 1 only for cores whose nodulated plants were dominated by the *nodA* allele corresponding to pSym type 3 (greater than 50% of *nodA* reads) and a 0 for all other cores, including non-nodulated cores. This nesting ensured pSym type and nodulation status were independent. These two predictors therefore correspond to sequential biological events (nodulation, and then pSym quality conditional on nodulation). Additionally, we included ASV-E relative abundance directly into this model, to allow for both indirect and direct ecological paths. All non-binary variables were Z-score standardized. We report standard effect sizes from both the ecological path (indirect * direct) as well as the evolutionary path. To understand if the magnitude of effect sizes was significantly different, we bootstrapped with 2000 replicates and report the 95^th^ percentile confidence interval and the two-sided *p* value.

*Generation of nodA phylogeny and stacked bar plot:* Each of the nodA ASVs which passed relative abundance and presence/absence filtering, along with references from our previous work classifying major pSym types 1-3, as well as three unique pSym types, were trimmed to the same length and aligned using MAFFT^S15^ on the local pair settings with 1000 iterations on 30 threads. We used IQ-TREE2^S16^ to infer phylogenetic relationships. The aligned sequences were analyzed using 1000 ultrafast bootstrap replicates (-B 1000) to assess branch support. Standard settings were employed; IQ-TREE2 automatically selected the best-fit substitution model. This tree was visualized in iTOL. As shown on the tree, trimming to align resulted in two ASVs, one very common and one rare, which have one SNP towards the end of the sequence, to be grouped into one ASV for Type 1. All analyses on the Type 1 pSym were done on the sum total reads of these two ASVs. To visualize pSym abundance change between nitrogen and control environments, a stacked bar plot of the major four pSym alleles (comprising over 98% of reads) was computed.

Plant biomass differences between N-addition and control were calculated using a Type 3 ANOVA on all three harvests, not considering uninoculated controls. Both field treatment and plant species were considered. Data are pooled across plant species because both exhibited similar responses to field treatment (no significant interaction

*Rhizobium genus Venn diagrams and stacked bar:* Venn diagrams of *Rhizobium* ASV presence-absence per compartment were computed using eulerr^S21^. An ASV had to be present in at least one sample to be considered present. Relative abundance of *Rhizobium* ASVs were visualized between compartments in a stacked bar plot. All ASVs with a relative abundance under 1% within a sample were grouped into “Other”. Relative abundance was normalized to 100%.

**R package details:** All analyses were performed in R 4.5.1 (2025-6-13). A complete list of packages and versions is available on Zenodo (10.5281/zenodo.17410936).

Figures 1, 2A, C, and D, 3, 5, and 6 A and C were made using ggplot. Figure 2B was visualized in iTOL. Figure 4 was visualized in iTOL. Figure 6B was made using euler. Supplementary figures 1, 3, 4, 5, and 6 were made using ggplot. Supplementary figure 2 was made using Vegan. Relativized molecular data (vegan::decostand, method “total”, in the vegan package^22^) were used for all analyses except for DESeq2^19^, which only accepts raw data. Data were pseudocounted for use in DESeq2, with 1 read being added to each *Rhizobium* ASV before modeling, to avoid all zero rows and allow DESeq2 to compute size factors within the *Rhizobium* subset^23,24^.

**Large language model versions:**

ChatGPT was used prior to January 2026, and Claude was used post January 2026. The versions of ChatGPT used were GPT-4o, o1, 4.5, o3, 5, and 5.4. The versions of Claude used were Opus 4.6, Opus 4.6 Extended, Opus 4.7, and Opus 4.7 Extended.

**Data availability:** All data is available on Zenodo and the NCBI SRA. All raw plant data: 10.5281/zenodo.17410894. Processed, unfiltered sequence files: Zenodo 10.5281/zenodo.17345394. All raw sequencing files are available in the NCBI SRA (BioProject PRJNA1338501). All scripts and code, as well as filtered sequencing data: 10.5281/zenodo.17410936.

**Supplementary Figures**

**Supplementary Figure 1:** **Absolute vs Relative Abundance of *Rhizobium* and ten randomly selected genera. A: Absolute and relative abundance are highly correlated in bulk soil for genus *Rhizobium.*** Spearman’s correlation of ten samples sequenced with a ZymoBiomics spike-in. Absolute cell counts calculated using a custom R script and correlated with relative abundance. **B: Absolute and relative abundance of ten random genera are highly correlated in bulk soil.** The same script was employed on ten randomly selected genera, to correlate absolute cell counts with relative abundance. Some low abundance genera include zeroes which can inflate apparent correlations; these are shown for completeness and to indicate general trends across taxa.

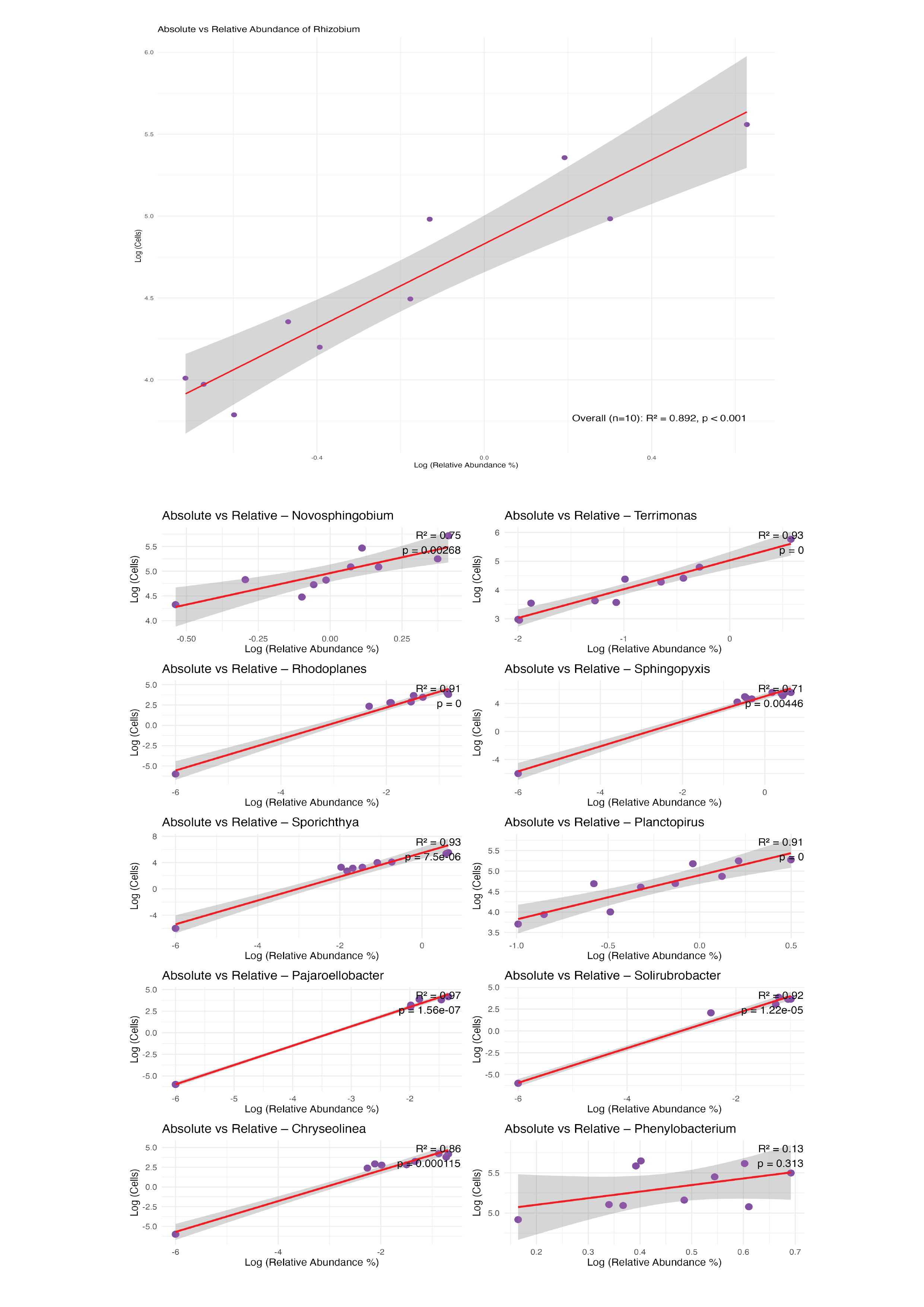

**Supplementary Figure 2: Rarefaction curves of bacterial community and genus *Rhizobium.* Top**: Filtering out low abundance ASVs still leaves the majority of meaningful diversity. Rarefaction curve of 303 samples. **Bottom**: Rhizobia remain diverse in all four compartments (source soil, bulk soil, rhizosphere, and nodules) when low-abundance taxa are filtered out. For both figures, ASVs were retained that were 1) in more than one sample, 2) had greater than 100 reads in at least one sample they were present in and 3) Were not found to be chimeric.

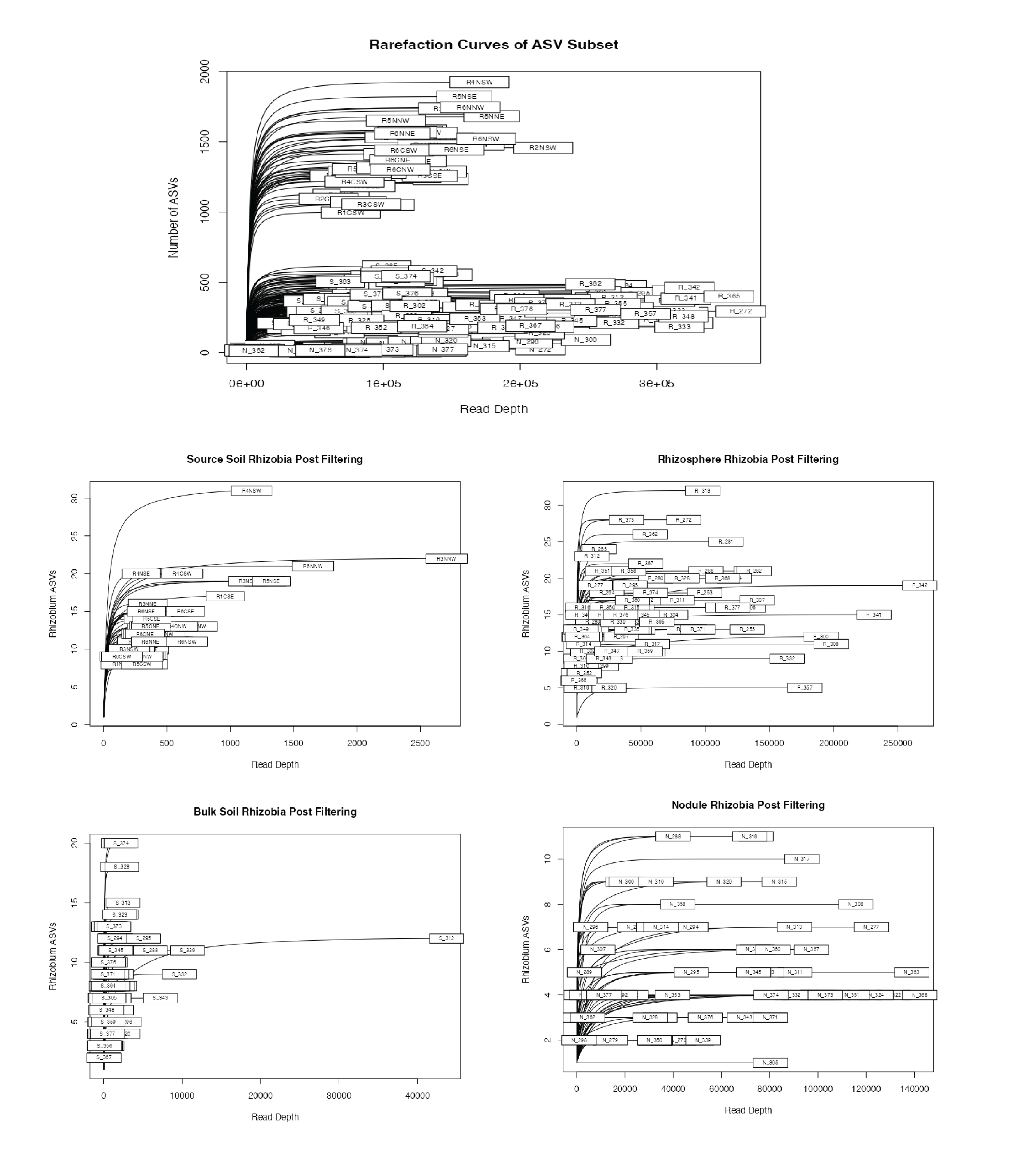

**Supplementary Figure 3:** **Distance-based redundancy analysis of the whole microbial community identified by compartment and nitrogen treatment.** Values computed from Bray-Curtis dissimilarity (full model in Supp Methods). Compartments are different colors, control plants are light shades and nitrogen plants are dark shades. Relative abundance data was used. Source soil appears linearly distributed due to a large compositional difference from the plant-associated samples. Axes show proportion of constrained variance explained.

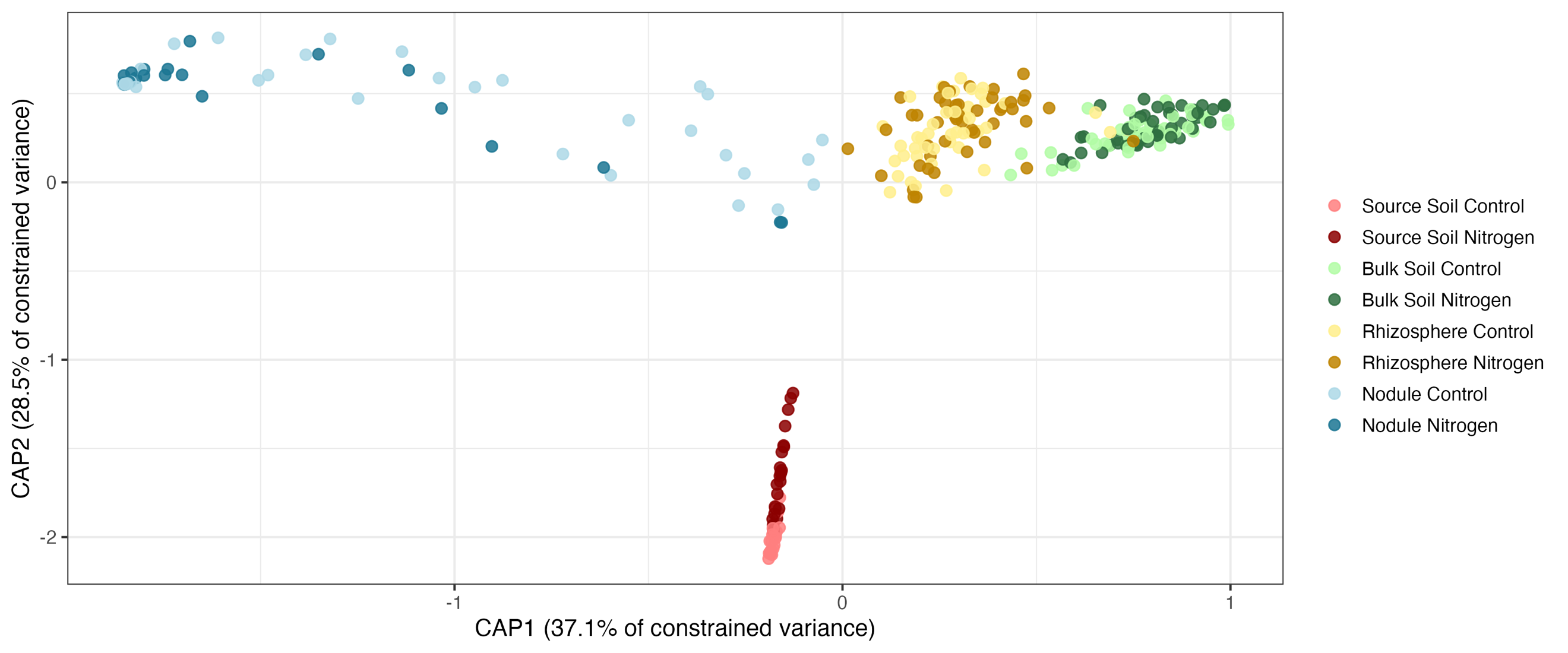

**Supplementary Figure 4: ~20% of nodule samples are dominated by genera other than *Rhizobium.*** Stacked bar plot of relative abundance of genera within nodule samples. Genera are colored, genera comprising less than 1% of reads are grouped into “Other”. Dominance is defined as whether or not a genus/genera other than *Rhizobium* comprises greater than 50% of relative abundance within a particular nodule sample.

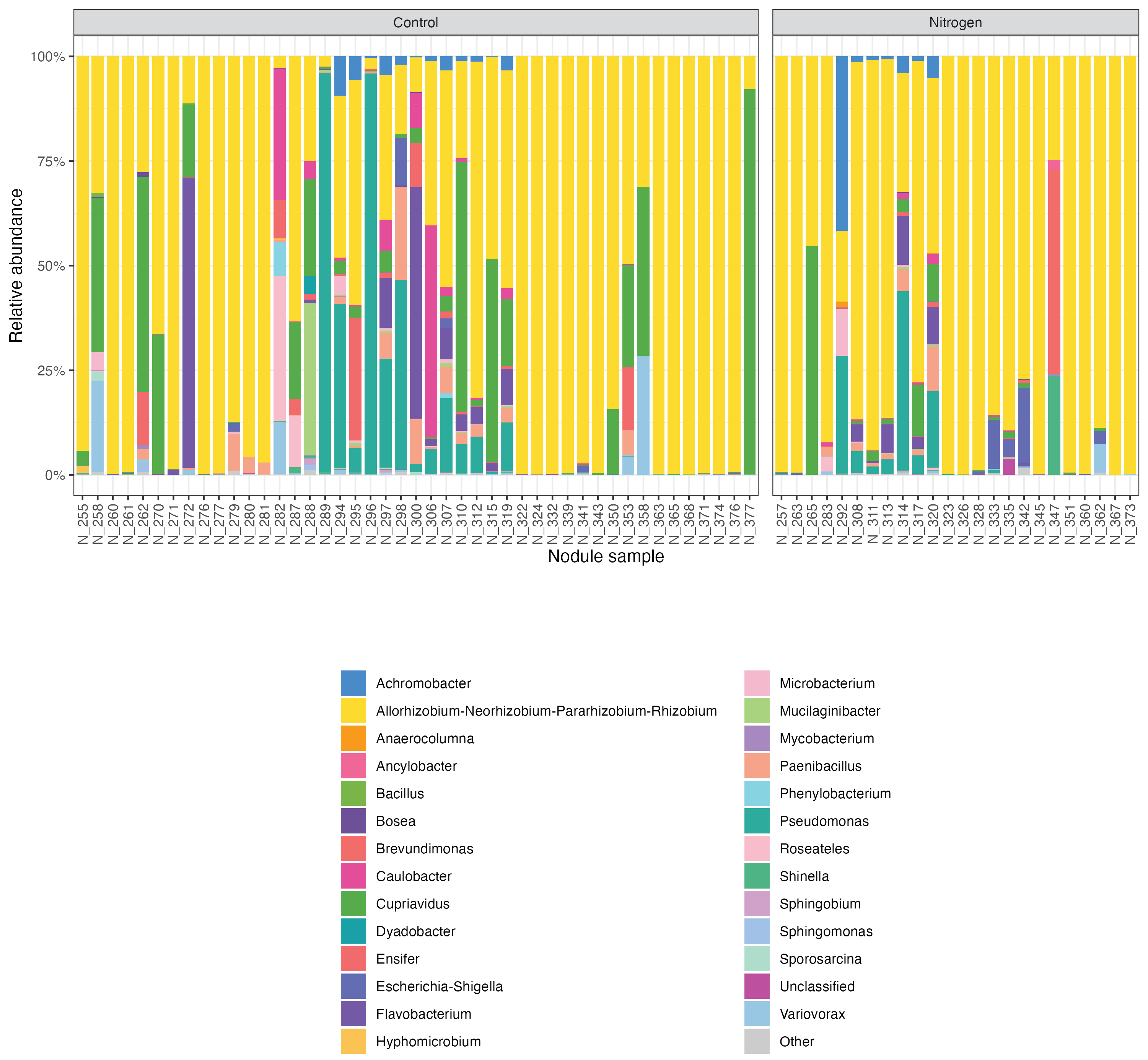

**Supplementary Figure 5: *ANPR* genera respond to compartment and nitrogen treatment.** **Distance-based redundancy analysis of ANPR ASVs identified by compartment and nitrogen treatment.** Values computed from Bray-Curtis dissimilarity (full model in Supp Methods). Compartments are different colors, control plants are light shades and nitrogen plants are dark shades. Relative abundance data was used.

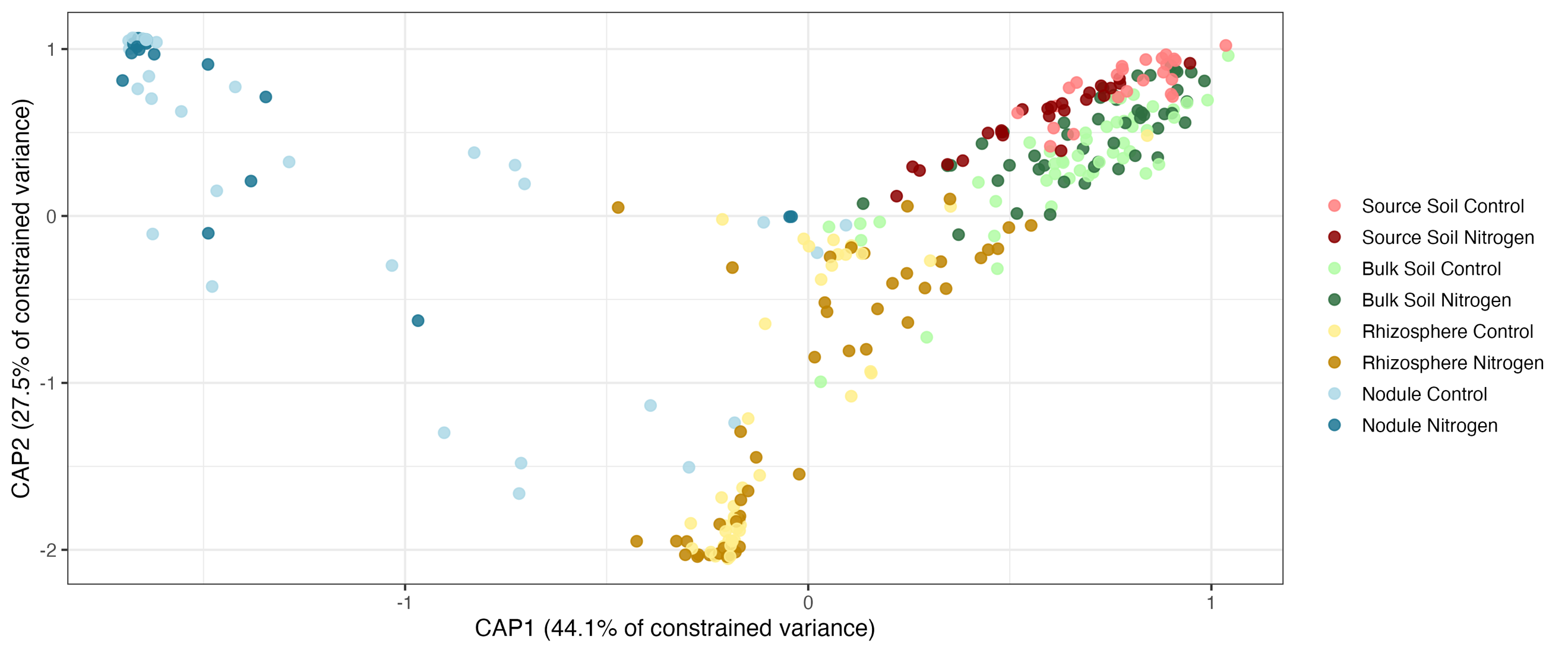

**Supplementary Figure 6: *Rhizobium* ASVs respond to compartment and nitrogen treatment.** **Distance-based redundancy analysis of Rhizobium ASVs identified by compartment and nitrogen treatment.** Values computed from Bray-Curtis dissimilarity (full model in Supp Methods). Compartments are different colors, control plants are light shades and nitrogen plants are dark shades. Relative abundance data was used.

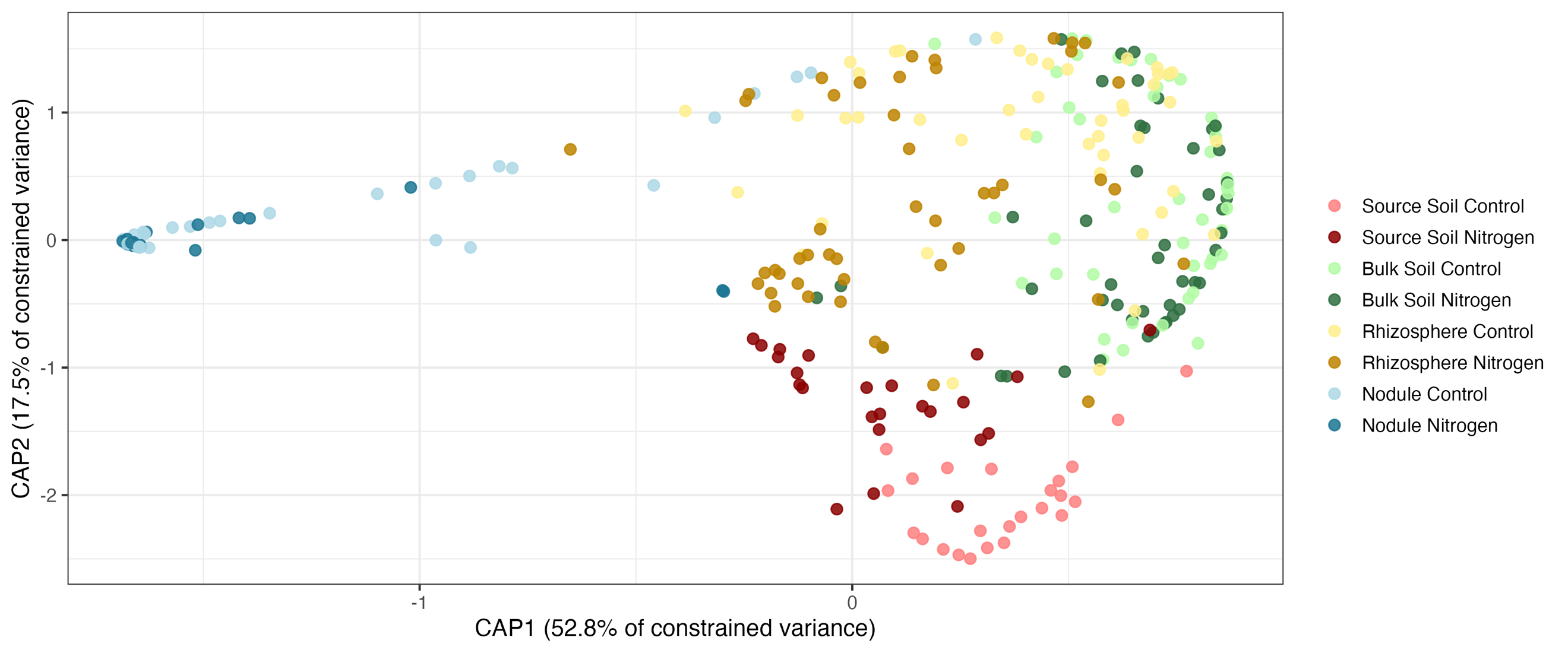

**Supplementary Figure 7: Biomass of plants inoculated with autoclave-killed (sterile) soil does not differ between N-addition and control.** N = 48 across three harvests. *P* = 0.8365 determined by Wilcoxon rank sum test.

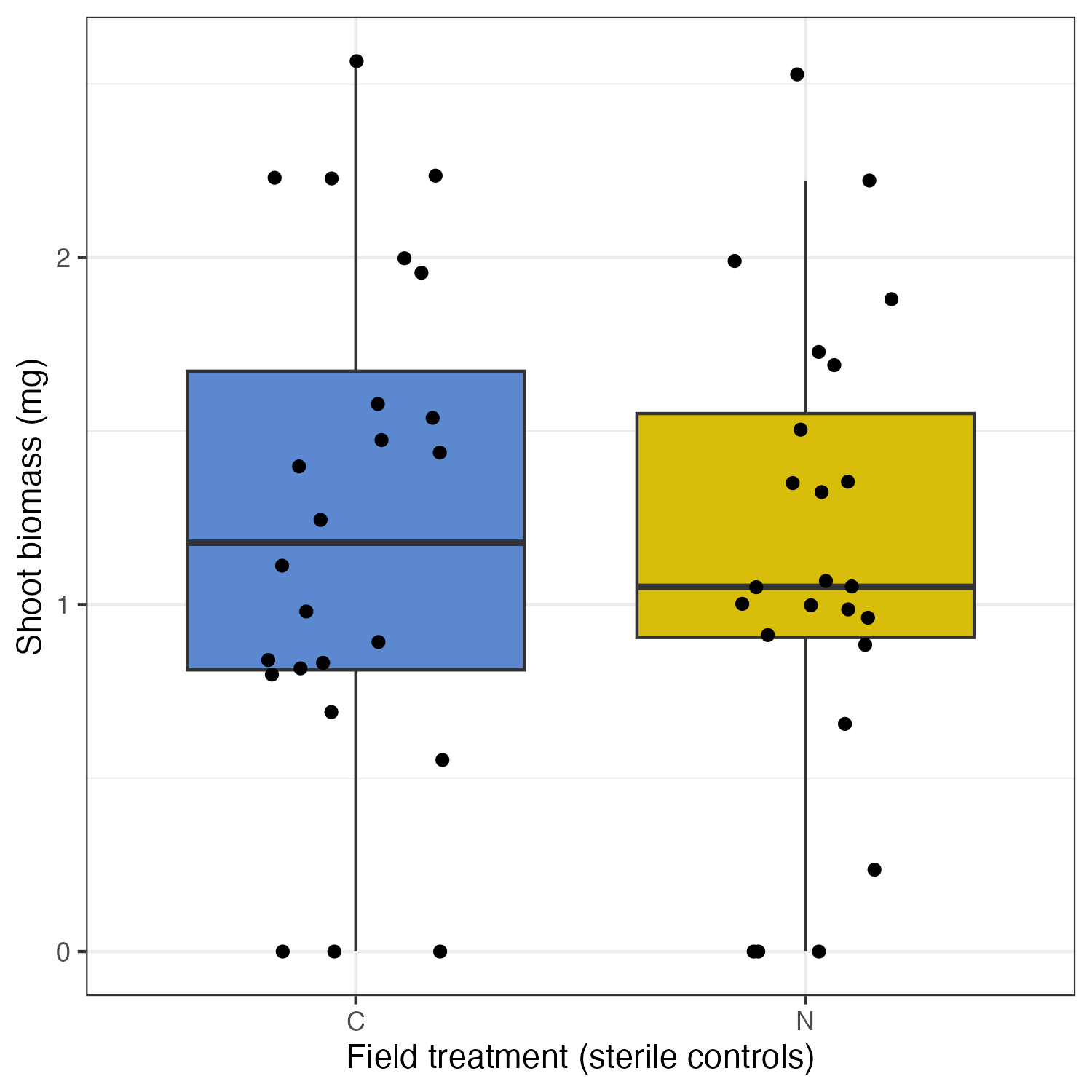

**Supplementary Tables**

**Supplementary Table 1: The whole microbial community responds strongly to compartment and nitrogen. A**: perMANOVA of the whole microbial community considering compartment, nitrogen treatment, plant species (*Trifolium hybridum* or *repens*), and plot (as a stratified factor). **B**: Compartment-specific perMANOVAs testing the difference between nitrogen and control environments.

**A:**

| **Factor** | **Df** | **SumOfSqs** | **R^2^** | **F** | **Pr(>F)** |
| --- | --- | --- | --- | --- | --- |
| Nitrogen_treat | 1 | 1.547 | 0.012 | 5.862 | **0.001** |
| Compartment | 3 | 40.903 | 0.324 | 51.651 | **0.001** |
| Plant_Spp | 1 | 1.011 | 0.008 | 3.830 | **0.001** |
| Nitrogen_treat:Compartment | 3 | 4.675 | 0.037 | 5.904 | **0.001** |
| Nitrogen_treat:Plant_Spp | 1 | 0.349 | 0.003 | 1.322 | 0.160 |
| Compartment:Plant_Spp | 2 | 0.977 | 0.008 | 1.850 | **0.008** |
| Nitrogen_treat:Compartment:Plant_Spp | 2 | 0.599 | 0.005 | 1.135 | 0.247 |
| Residual | 289 | 76.288 | 0.604 |  |  |
| Total | 302 | 126.350 | 1.000 |  |  |

**B:**

| **Compartment** | **source_soil** | **bulk_soil** | **rhizosphere** | **nodule** |
| --- | --- | --- | --- | --- |
| **R^2^** | 0.318 | 0.034 | 0.035 | 0.023 |
| **p** | 0.001 | 0.001 | 0.002 | 0.149 |

**Supplementary Table 2: Linear mixed models of ASV-E relative abundance separated by compartment.** Plot was included as a random variable.

| **Compartment** | **Factor** | **Sum Sq** | **Mean Sq** | **NumDF** | **DenDF** | **F value** | **Pr(>F)** |
| --- | --- | --- | --- | --- | --- | --- | --- |
| source_soil | Nitrogen_treat | 9.51e-07 | 9.51e-07 | 1 | 41 | 22.85 | **2.27e-05** |
| bulk_soil | Nitrogen_treat | 0.00015 | 0.00015 | 1 | 87.96 | 6.28 | **0.0141** |
| rhizosphere | Nitrogen_treat | 3.56e-05 | 3.56e-05 | 1 | 85.50 | 0.45 | 0.50 |
| nodule | Nitrogen_treat | 0.094 | 0.094 | 1 | 67 | 1.40 | 0.24 |

**Supplementary Table 3: SEM coefficients + standardized effects.**

**A: Path coefficients**

| **Response** | **Predictor** | **Estimate** | **Std. Error** | **DF** | **Crit. Value** | **P-value** | **Std. Estimate** |
| --- | --- | --- | --- | --- | --- | --- | --- |
| nod_pa | gse_relabun | 1.95 | 0.939 | 46 | 2.08 | **0.0379** | 0.732 |
| mean_biomass | nod_pa | 1.28 | 0.382 | 44 | 3.35 | **0.00170** | 0.457 |
| mean_biomass | psym_type3 | -0.793 | 0.426 | 44 | -1.86 | 0.0692 | -0.245 |
| mean_biomass | gse_relabun | 0.106 | 0.136 | 44 | 0.777 | 0.441 | 0.106 |

**B: standardized effects**

| **Effect** | **Std. value** |
| --- | --- |
| Ecological indirect (gsE → nod → biomass) | 0.334 |
| Ecological direct (gsE → biomass) | 0.106 |
| Ecological total (indirect + direct) | 0.440 |
| Evolutionary direct (pSym → biomass) | -0.245 |
| Ratio (\|eco total\| / \|evo\|) | 1.80 |
| Difference (\|eco\| − \|evo\|) | 0.190 |
| 95% CI of difference | -0.207 to 0.570 |
| Two-sided p-value | 0.333 |

**Supplementary Table 4: Type III ANOVAs of A) Relative abundance, B) Shannon diversity, C) Evenness and D) Richness of the genus *Rhizobium.*** Field treatment, compartment, and the interaction of nitrogen and compartment were fixed predictor variables, with plot as a random factor. Significance annotations in the corresponding main-text fig. (Fig. 5) are based on Tukey’s post-hoc tests. Because source soil samples were not part of the greenhouse design/had no plant species associated, we computed separate models that removed these samples and included plant species as a fixed effect. Inclusion of plant species did not alter relative abundance, richness or evenness patterns and only modestly affected Shannon diversity, so results are not further reported.

| **Factor** | **Sum Sq** | **Mean Sq** | **NumDF** | **DenDF** | **F** | **Pr(>F)** |
| --- | --- | --- | --- | --- | --- | --- |
| ***A: Relative abundance*** | | | | | | |
| Nitrogen_treat | 0.088 | 0.088 | 1 | 295.00 | 6.596 | **0.011** |
| Compartment | 12.031 | 4.010 | 3 | 295.00 | 301.287 | **<0.001** |
| Nitrogen_treat:Compartment | 0.212 | 0.071 | 3 | 295.00 | 5.308 | **0.001** |
| ***B: Shannon diversity*** | | | | | | |
| Nitrogen_treat | 0.159 | 0.159 | 1 | 291.94 | 1.192 | 0.276 |
| Compartment | 56.632 | 18.877 | 3 | 291.30 | 141.807 | **<0.001** |
| Nitrogen_treat:Compartment | 0.559 | 0.186 | 3 | 291.10 | 1.400 | 0.243 |
| ***C: Richness*** | | | | | | |
| Nitrogen_treat | 0.850 | 0.851 | 1 | 292.91 | 0.358 | 0.550 |
| Compartment | 532.400 | 177.466 | 3 | 291.30 | 74.701 | **<0.001** |
| Nitrogen_treat:Compartment | 1.990 | 0.665 | 3 | 291.65 | 0.280 | 0.840 |
| ***D: Evenness*** | | | | | | |
| Nitrogen_treat | 0.004 | 0.004 | 1 | 242.59 | 0.070 | 0.791 |
| Compartment | 15.725 | 5.242 | 3 | 241.88 | 104.102 | **<0.001** |
| Nitrogen_treat:Compartment | 0.493 | 0.164 | 3 | 241.61 | 3.263 | **0.022** |

**Supplementary Table 5: The *ANPR* genera respond strongly to nitrogen, compartment, and their interaction.** A: perMANOVA of the ANPR ASVs considering compartment, nitrogen treatment, plant species, and plot (as a stratified factor). B: Compartment-specific perMANOVAs testing the difference between nitrogen and control environments.

**A:**

| **Factor** | **Df** | **SumOfSqs** | **R^2^** | **F** | **Pr(>F)** |
| --- | --- | --- | --- | --- | --- |
| Nitrogen_treat | 1 | 1.861 | 0.015 | 6.964 | **0.001** |
| Compartment | 3 | 40.425 | 0.324 | 50.428 | **0.001** |
| Plant_Spp | 1 | 0.311 | 0.002 | 1.162 | 0.279 |
| Nitrogen_treat:Compartment | 3 | 3.643 | 0.029 | 4.544 | **0.001** |
| Nitrogen_treat:Plant_Spp | 1 | 0.385 | 0.003 | 1.442 | 0.143 |
| Compartment:Plant_Spp | 2 | 0.530 | 0.004 | 0.993 | 0.438 |
| Nitrogen_treat:Compartment:Plant_Spp | 2 | 0.509 | 0.004 | 0.953 | 0.484 |
| Residual | 289 | 77.226 | 0.618 |  |  |
| Total | 302 | 124.890 | 1.000 |  |  |

**B:**

| **Compartment** | **source_soil** | **bulk_soil** | **rhizosphere** | **nodule** |
| --- | --- | --- | --- | --- |
| **R^2^** | 0.193 | 0.051 | 0.036 | 0.029 |
| **p** | 0.001 | 0.001 | 0.007 | 0.097 |

**Supplementary Table 6: The *Rhizobium* genus responds strongly to nitrogen, compartment, and their interaction.** A: perMANOVA of the *Rhizobium* ASVs considering compartment, nitrogen treatment, plant species, and plot (as a stratified factor). B: Compartment-specific perMANOVAs testing the difference between nitrogen and control environments.

**A:**

| **Factor** | **Df** | **SumOfSqs** | **R^2^** | **F** | **Pr(>F)** |
| --- | --- | --- | --- | --- | --- |
| Nitrogen_treat | 1 | 3.946 | 0.035 | 17.641 | **0.001** |
| Compartment | 3 | 37.893 | 0.336 | 56.463 | **0.001** |
| Plant_Spp | 1 | 0.347 | 0.003 | 1.548 | 0.121 |
| Nitrogen_treat:Compartment | 3 | 4.758 | 0.042 | 7.090 | **0.001** |
| Nitrogen_treat:Plant_Spp | 1 | 0.224 | 0.002 | 1.001 | 0.396 |
| Compartment:Plant_Spp | 2 | 0.658 | 0.006 | 1.470 | 0.104 |
| Nitrogen_treat:Compartment:Plant_Spp | 2 | 0.368 | 0.003 | 0.823 | 0.640 |
| Residual | 289 | 64.650 | 0.573 |  |  |
| Total | 302 | 112.843 | 1.000 |  |  |

**B:**

| **Compartment** | **source_soil** | **bulk_soil** | **rhizosphere** | **nodule** |
| --- | --- | --- | --- | --- |
| **R^2^** | 0.263 | 0.053 | 0.125 | 0.036 |
| **p** | 0.001 | 0.002 | 0.001 | 0.076 |

**References Shared with Main Text:**

38.	Weese, D. J., Heath, K. D., Dentinger, B. T. M. & Lau, J. A. Long-term nitrogen addition causes the evolution of less-cooperative mutualists. *Evolution* **69**, 631–642 (2015).

42.	Naranjo-Robayo, N. *et al.* Updated taxonomy of the family Rhizobiaceae with proposals for 10 novel genera and 35 novel combinations. 2025.11.20.689363 Preprint at https://doi.org/10.1101/2025.11.20.689363 (2025).

43.	Young, J. P. W. *et al.* Defining the Rhizobium leguminosarum Species Complex. *Genes* **12**, 111 (2021).

51.	Huberty, L. E., Gross, K. L. & Miller, C. J. Effects of nitrogen addition on successional dynamics and species diversity in Michigan old-fields. *J. Ecol.* **86**, 794–803 (1998).

52.	Dickson, T. L. & Gross, K. L. Plant community responses to long-term fertilization: changes in functional group abundance drive changes in species richness. *Oecologia* **173**, 1513–1520 (2013).

53.	Kay Gross & Jennifer Lau. Plant Community and Ecosystem Responses to Long-term Fertilization and Disturbance at the Kellogg Biological Station. Environmental Data Initiative https://doi.org/https://doi.org/10.6073/pasta/ea22c735ddfe17c595ccca978a87d109.

**Supplementary References**

1.	Somasegaran, P. & Hoben, H. J. *Handbook for Rhizobia*. (Springer, New York, NY, 1994). doi:10.1007/978-1-4613-8375-8.

2.	Sachs, J. L., Kembel, S. W., Lau, A. H. & Simms, E. L. In Situ Phylogenetic Structure and Diversity of Wild Bradyrhizobium Communities. *Appl. Environ. Microbiol.* **75**, 4727–4735 (2009).

3.	Batstone, R. T., O’Brien, A. M., Harrison, T. L. & Frederickson, M. E. Experimental evolution makes microbes more cooperative with their local host genotype. *Science* **370**, 476–478 (2020).

4.	Haukka, K., Lindström, K. & Young, J. P. W. Three Phylogenetic Groups of nodA and nifHGenes in Sinorhizobium and Mesorhizobium Isolates from Leguminous Trees Growing in Africa and Latin America. *Appl. Environ. Microbiol.* **64**, 419–426 (1998).

5.	Callahan, B. J. *et al.* DADA2: High resolution sample inference from Illumina amplicon data. *Nat. Methods* **13**, 581–583 (2016).

6.	Lennard, K. *et al.* TADA: Beta v0.5 release. https://doi.org/10.5281/zenodo.4208836 (2020).

7.	Blum, M. *et al.* InterPro: the protein sequence classification resource in 2025. *Nucleic Acids Res.* **53**, D444–D456 (2025).

8.	Lan, Y., Wang, Q., Cole, J. R. & Rosen, G. L. Using the RDP Classifier to Predict Taxonomic Novelty and Reduce the Search Space for Finding Novel Organisms. *PLOS ONE* **7**, e32491 (2012).

9.	Quast, C. *et al.* The SILVA ribosomal RNA gene database project: improved data processing and web-based tools. *Nucleic Acids Res.* **41**, D590-596 (2013).

10.	McLaren, M. Silva Reference Database. Zenodo https://doi.org/10.5281/zenodo.4587946 (2021).

11.	Steinegger, M. & Söding, J. MMseqs2 enables sensitive protein sequence searching for the analysis of massive data sets. *Nat. Biotechnol.* **35**, 1026–1028 (2017).

12.	McMurdie, P. J. & Holmes, S. Waste Not, Want Not: Why Rarefying Microbiome Data Is Inadmissible. *PLOS Comput. Biol.* **10**, e1003531 (2014).

13.	Zhou, R., Ng, S. K., Sung, J. J. Y., Goh, W. W. B. & Wong, S. H. Data pre-processing for analyzing microbiome data – A mini review. *Comput. Struct. Biotechnol. J.* **21**, 4804–4815 (2023).

14.	Tan, C. C. S. *et al.* No evidence for a common blood microbiome based on a population study of 9,770 healthy humans. *Nat. Microbiol.* **8**, 973–985 (2023).

15.	Katoh, K. & Standley, D. M. MAFFT Multiple Sequence Alignment Software Version 7: Improvements in Performance and Usability. *Mol. Biol. Evol.* **30**, 772–780 (2013).

16.	Minh, B. Q. *et al.* IQ-TREE 2: New Models and Efficient Methods for Phylogenetic Inference in the Genomic Era. *Mol. Biol. Evol.* **37**, 1530–1534 (2020).

17.	Seemann, T. Barrnap. (2025).

18.	Kuznetsova, A., Brockhoff, P. B. & Christensen, R. H. B. lmerTest Package: Tests in Linear Mixed Effects Models. *J. Stat. Softw.* **82**, 1–26 (2017).

19.	Love, M. I., Huber, W. & Anders, S. Moderated estimation of fold change and dispersion for RNA-seq data with DESeq2. *Genome Biol.* **15**, 550 (2014).

20.	Lefcheck, J. S. piecewiseSEM: Piecewise structural equation modelling in r for ecology, evolution, and systematics. *Methods Ecol. Evol.* **7**, 573–579 (2016).

21.	Larsson, J. & Gustafsson, P. A Case Study in Fitting Area-Proportional {{Euler}} Diagrams with Ellipses Using eulerr. *Proceedings of international workshop on set visualization and reasoning 2018* (2018).

22.	vegan: an R package for community ecologists. https://vegandevs.github.io/vegan/.

23.	Fan, X. *et al.* Drinking alcohol is associated with variation in the human oral microbiome in a large study of American adults. *Microbiome* **6**, 59 (2018).

24.	Peters, B. A. *et al.* A taxonomic signature of obesity in a large study of American adults. *Sci. Rep.* **8**, 9749 (2018).
